# Drug Discovery in Low Data Regimes: Leveraging a Computational Pipeline for the Discovery of Novel SARS-CoV-2 Nsp14-MTase Inhibitors

**DOI:** 10.1101/2023.10.03.560722

**Authors:** AkshatKumar Nigam, Matthew F. D. Hurley, Fengling Li, Eva Konkoľová, Martin Klíma, Jana Trylčová, Robert Pollice, Süleyman Selim Çinaroğlu, Roni Levin-Konigsberg, Jasemine Handjaya, Matthieu Schapira, Irene Chau, Sumera Perveen, Ho-Leung Ng, H. Ümit Kaniskan, Yulin Han, Sukrit Singh, Christoph Gorgulla, Anshul Kundaje, Jian Jin, Vincent A. Voelz, Jan Weber, Radim Nencka, Evzen Boura, Masoud Vedadi, Alán Aspuru-Guzik

**Affiliations:** Department of Computer Science, Stanford University; Department of Genetics, Stanford University; Department of Chemistry, Temple University, Philadelphia, PA 19122, USA; Structural Genomics Consortium, University of Toronto, Toronto, Ontario M5G 1L7, Canada; Institute of Organic Chemistry and Biochemistry of the Czech Academy of Sciences, Prague, Czech Republic; Chemical Physics Theory Group, Department of Chemistry, University of Toronto, 80 St. George St, Toronto, Ontario M5S 3H6, Canada; Department of Computer Science, University of Toronto, 40 St. George St, Toronto, Ontario M5S 2E4, Canada; Stratingh Institute for Chemistry, University of Groningen, The Netherlands; Structural Bioinformatics and Computational Biochemistry, Department of Biochemistry, University of Oxford, South Parks Road, Oxford, OX1 3QU, UK; Department of Pharmacology and Toxicology, University of Toronto, Toronto, Ontario M5S 1A8, Canada; Department of Biochemistry and Molecular Biophysics, Kansas State University, Manhattan, KS 66506, USA; Department of Pharmacological Sciences and Oncological Sciences, Mount Sinai Center for Therapeutics Discovery, Tisch Cancer Institute, Ichan School of Medicine at Mount Sinai, New York, NY, USA; Computational and Systems Biology Program, Memorial Sloan Kettering Cancer Center; St. Jude Children’s Research Hospital, Department of Structural Biology, Memphis, TN, USA; Department of Physics, Faculty of Arts and Sciences, Harvard University, Cambridge, USA; Department of Cancer Biology, Dana-Farber Cancer Institute, Boston, USA; QBI COVID-19 Research Group (QCRG), San Francisco, CA, USA; Drug Discovery Program, Ontario Institute for Cancer Research, Toronto, Ontario, Canada; Department of Chemical Engineering & Applied Chemistry, University of Toronto, Canada; Department of Materials Science & Engineering, University of Toronto, Canada; Vector Institute for Artificial Intelligence, Toronto, Canada; Canadian Institute for Advanced Research (CIFAR), Toronto, ON, Canada; Acceleration Consortium, University of Toronto, Toronto, ON, Canada

## Abstract

The COVID-19 pandemic, caused by the SARS-CoV-2 virus, has led to significant global morbidity and mortality. A crucial viral protein, the non-structural protein 14 (nsp14), catalyzes the methylation of viral RNA and plays a critical role in viral genome replication and transcription. Due to the low mutation rate in the nsp region among various SARS-CoV-2 variants, nsp14 has emerged as a promising therapeutic target. However, discovering potential inhibitors remains a challenge. In this work, we introduce a computational pipeline for the rapid and efficient identification of potential nsp14 inhibitors by leveraging virtual screening and the NCI open compound collection, which contains 250,000 freely available molecules for researchers worldwide. The introduced pipeline provides a cost-effective and efficient approach for early-stage drug discovery by allowing researchers to evaluate promising molecules without incurring synthesis expenses. Our pipeline successfully identified seven promising candidates after experimentally validating only 40 compounds. Notably, we discovered NSC620333, a compound that exhibits a strong binding affinity to nsp14 with a dissociation constant of 427 ± 84 nM. In addition, we gained new insights into the structure and function of this protein through molecular dynamics simulations. We identified new conformational states of the protein and determined that residues Phe367, Tyr368, and Gln354 within the binding pocket serve as stabilizing residues for novel ligand interactions. We also found that metal coordination complexes are crucial for the overall function of the binding pocket. Lastly, we present the solved crystal structure of the nsp14-MTase complexed with SS148 (PDB:8BWU), a potent inhibitor of methyltransferase activity at the nanomolar level (IC_50_ value of 70 ± 6 nM). Our computational pipeline accurately predicted the binding pose of SS148, demonstrating its effectiveness and potential in accelerating drug discovery efforts against SARS-CoV-2 and other emerging viruses.

## I. INTRODUCTION

The COVID-19 pandemic, caused by the Severe Acute Respiratory Syndrome Coronavirus 2 (SARS-CoV-2), has had a profound impact on global health, economies, and daily life. SARS-CoV-2 is a single-stranded, positive-sense RNA virus belonging to lineage B of the genus Beta-coronavirus in the Coronaviridae family [1]. Since its emergence in December 2019, it has led to a staggering number of infections and deaths world-wide, overwhelming healthcare systems and prompting unprecedented public health measures [2]. While the development of vaccines has been a significant step in combating the virus, the continued emergence of new variants highlights the ongoing need for effective antiviral treatments. Such treatments can play a crucial role in reducing the severity and impact of infections and informing future pandemic policy decisions [3]. The SARS-CoV-2 genome is notably large, containing nearly 30,000 base pairs and encoding 16 nonstructural proteins (nsps) that play vital roles in viral replication and transcription [4]. Because of this, nsps represent potential targets for therapeutics, and the availability of their structural data has facilitated the rapid development of antivirals [5]. One such target is non-structural protein 14 (nsp14), which has dual functions as an exonuclease (ExoN) and a methyltransferase (MTase). The ExoN function ensures replication fidelity through 3’-5’ proofreading, while the MTase function catalyzes N7-methylation of the guanosine triphosphate cap of the viral RNA, which is crucial for its stability and function [6, 7]. Due to its low mutation rate among different SARS-CoV-2 variants when compared to other viral proteins such as the spike protein [8], nsp14 is hypothesized to be essential for the function of the virus, and therefore it has emerged as a promising therapeutic target. This persistence increases the likelihood that inhibitors targeting nsp14 will remain effective against a broader range of evolving viral strains. Additionally, the amino acid sequence of SARS-CoV-2 nsp14 shows high homology with nsp14 from other coronaviruses, including SARS-CoV, MERS-CoV, and common human coronaviruses such as OC43, HKU1, 229E, and NL63 [8]. This sequence conservation suggests that inhibitors targeting nsp14 in SARS-CoV-2 could potentially also inhibit nsp14 MTase activity in other coronaviruses, which may have implications for the development of broad-spectrum antiviral drugs [9].

In this study, we developed a computational pipeline for rapid and efficient identification of potential nsp14 inhibitors. By leveraging virtual screening at various levels of accuracy and the NCI open compound collection [10, 11]—a database comprising 250,000 freely accessible molecules for researchers globally, we evaluated promising molecules without the expense and time typically associated with synthesis. This cost-effective and efficient strategy proves particularly valuable for early-stage hit identification, in cases where resources are constrained and time is of the essence. We utilized molecular docking [12, 13] and Molecular Mechanics Generalized Born Surface Area (MM/GBSA) calculations [14, 15] to gauge the strength of interaction between potential inhibitors and nsp14. Through these computational methodologies, we successfully identified a novel inhibitor for nsp14, with activity in the micromolar range, labelled NSC620333 (IC_50_ value of 5.3*±*1.0 *μM*), which was confirmed by orthogonal in vitro and in vivo assays. This inhibitor was unearthed through computational screening of the NCI open compound collection, with only the top 40 compounds undergoing experimental testing, highlighting the efficiency of our pipeline. Furthermore, to deepen our understanding of the protein structure and function, we conducted molecular dynamics (MD) simulations [16– 18] exceeding 5 microseconds. These simulations unveiled previously unknown conformational states of nsp14, offering valuable insights into its overall function. Within the MTase lateral cavity, we identified flexible residues— specifically Phe367, Tyr368, and Gln354—that form stabilizing interactions with potential ligands. Our simulations also revealed that the two alpha-helices of the MTase domain undergo a significant shift/bend, corresponding to the opening and closing of the lateral cavity. Importantly, these conformational shifts were not observed in the absence of metal coordination complexes, emphasizing their crucial role in stabilizing interactions. Our MD simulations also shed light on the interaction of nsp14 with nsp10, a key process in viral replication and transcription. Nsp10 has been demonstrated to stimulate the ExoN and MTase activities of nsp14, making it a critical cofactor for the functionality of the protein [19]. In summary, our research showcases an advanced computational pipeline, built upon the robust capabilities of VirtualFlow 2.0 [20], specifically the VirtualFlow Unity (VFU) module. This component not only accommodates a wide spectrum of docking programs, callable through a Python pipeline, but also allows scoring at diverse levels of precision, thereby facilitating the rapid and proficient detection of prospective nsp14 inhibitors. This is achieved by integrating virtual screening strategies and harnessing the expansive NCI open compound collection. Through this methodology, we discovered NSC620333, a compound exhibiting strong binding affinity to nsp14, and gained fresh insights into the structure and function of the protein. Furthermore, we report the crystal structure of SS148, a potent inhibitor (IC_50_ value of 70 ± 6 nM) of methyltransferase activity, in complex with the nsp14-MTase domain [21, 22]. Our pipeline accurately predicted the binding pose of SS148, demonstrating its efficacy and potential for expediting drug discovery efforts against SARS-CoV-2 and other emerging viruses. By targeting the nsp14 protein, we aspire to contribute to the development of effective antiviral therapies against SARS-CoV-2 and its variants, and other coronaviruses as well. Our computational pipeline offers a cost-effective and efficient strategy for early-stage drug discovery, potentially providing significant assistance to researchers battling the rapidly mutating virus responsible for COVID-19 and aiding in the response to future disease outbreaks.

## II. RESULTS

### A. Employing Molecular Docking for identifying nsp14 inhibitors

The widespread use of structure-based drug discovery techniques has proven instrumental in the search for small molecules that can specifically target macromolecules. [23] The efficacy of molecular docking and free energy simulation methods is particularly noteworthy in assessing the binding strength between proteins and ligands. [24] These methods enabled us to identify and prioritize the most promising candidates, thereby facilitating subsequent experimental testing. Our aim was to identify inhibitors of the SARS-CoV-2 nsp14 MTase activity. To achieve this, we performed a computational screening of the Developmental Therapeutics Program (DTP) Open Compound Collection. [10, 11] Comprising approximately 250,000 molecules synthesized and tested for potential efficacy against cancer and acquired immunodeficiency syndrome, this collection offers a considerable resource for researchers. Importantly, these molecules are readily accessible for academic researchers, with the possibility to request up to 40 samples on a monthly basis.

We employed the Glide [25] SP docking program to target the MTase binding site, as depicted in Figure I. This approach allowed us to evaluate all the molecules in the ligand library. To enhance the reliability of our findings, we rigorously examined the top 1,000 molecules— those displaying the lowest docking scores—using the Glide XP setting to increase precision. Subsequently, we selected the top 100 performing molecules for further consideration, factoring in aspects such as diversity (see Methods: Docking V A, compound availability, and docking scores. In an effort to further refine our selection and improve the accuracy of binding prediction, we conducted a Molecular Mechanics Generalized Born Surface Area (MM/GBSA) analysis [14, 15] (see Methods: MM/GBSA Calculation V B). This step assisted us in narrowing down our list to the top 40 compounds, which we subsequently obtained from the NCI. Supplementary Table S2 lists the selected compounds, along with their corresponding docking scores and Molecular Mechanics Generalized Born Surface Area (MM/GBSA) binding affinities. Notably, NSC620333 stood out, showcasing the most promising results, as demonstrated by its lowest docking and MM/GBSA scores. In Figure 2, we provide the docked poses and 2D interaction diagrams of the three well performing inhibitors. These 2D interaction diagrams highlight the fact that the majority of residues in the binding pocket are hydrophobic or non-polar. Detailed information regarding the docking procedure and protein preparation can be found in the Methods section (see Methods: Docking V A).

**FIG. 1.**
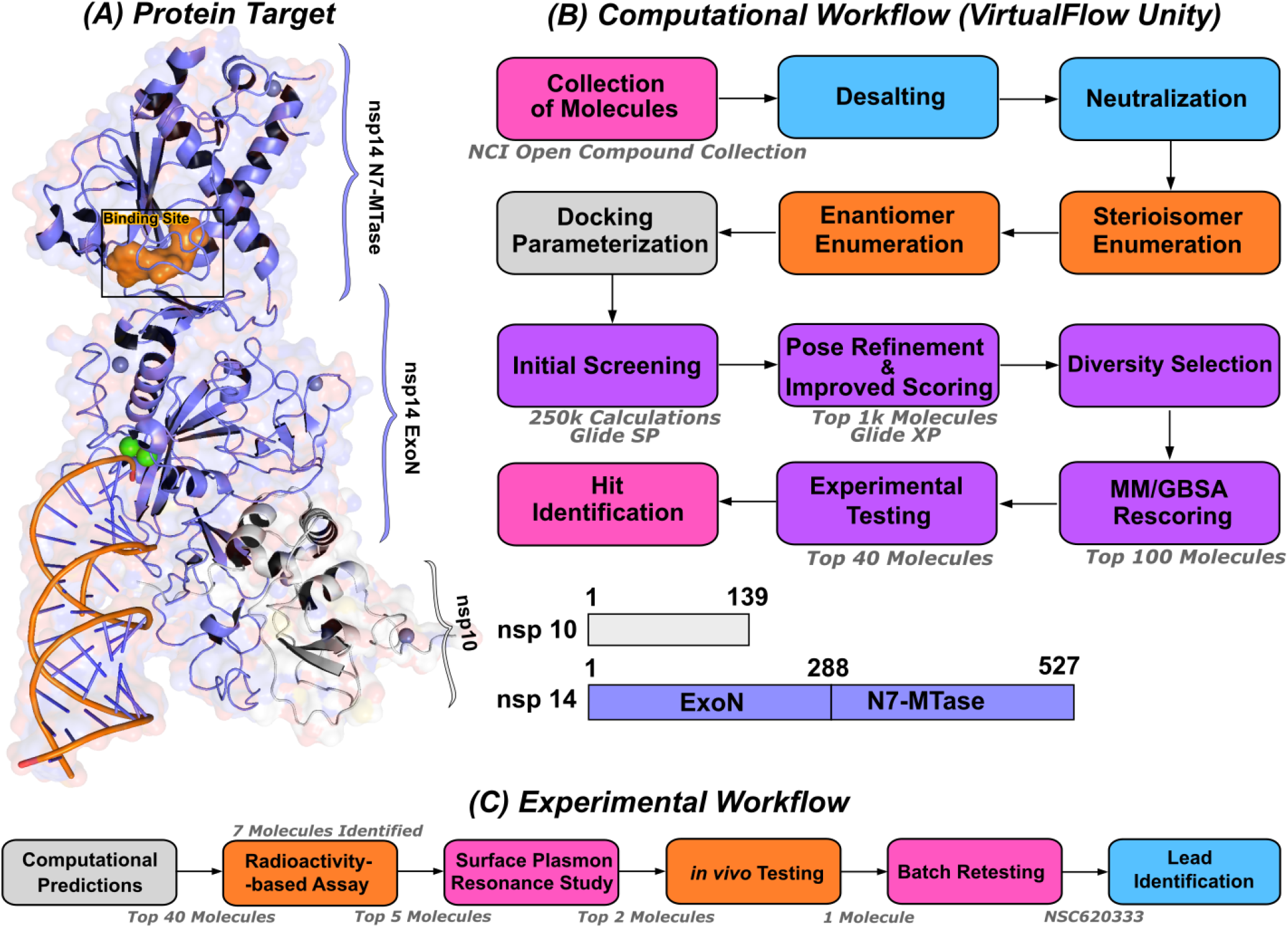
Computational pipeline for identification of lead compounds. **(A)** Depiction of the SARS-CoV-2 nsp14-MTase Protein Target and Computational Workflow. The protein target (PDB:7N0B), in complex with nsp10, is employed for the modeling process. The highlighted binding site (indicated in orange) within the MTase domain acts as the docking site for molecular interactions. **(B)** The computational workflow powered by VirtualFlow Unity (VFU), which processes SMILES strings from the NCI Open Compound Collections as input. The procedure includes desalting, neutralizing, and generating stereoisomers of the input molecule. Subsequently, all processed molecules from the database undergo docking onto the binding site using Glide-SP. The output docked compounds are ranked based on their docking scores, with the top 1,000 subjected to a more precise scoring algorithm, Glide-XP. A diverse subset of 100 molecules is chosen for an advanced binding energy estimation using Molecular Mechanics Generalized Born Surface Area (MM/GBSA). The top 40 of these compounds, ranked by their binding potential, are then selected for experimental evaluation. **(C)** The selected top-40 compounds are initially assessed using a radioactivity-based assay, yielding seven compounds with promising results. Subsequently, a detailed binding study is performed with the top-5 compounds utilizing surface plasmon resonance. The in-vivo evaluation follows, focusing on the top-2 compounds with SARS-CoV-2 infected Calu-3 cells. Ultimately, the compound identified as the most promising is re-tested after a custom synthesis process, leading not only to improved purity, but also confirming the identification of a lead compound.

**FIG. 2.**
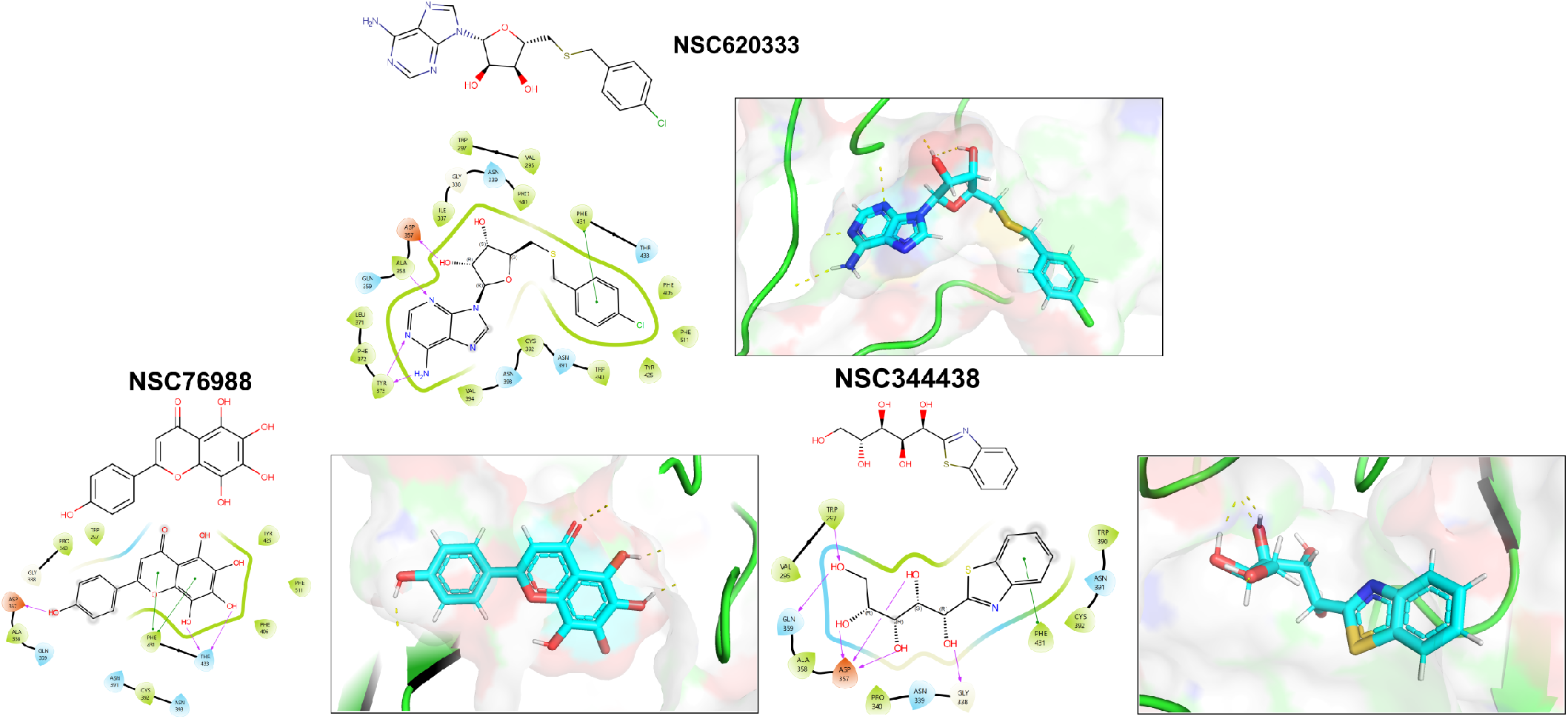
Interaction Analysis of Identified MTase Inhibitors. Presented are the 2D structural diagrams of experimentally validated compounds (top-3) found to inhibit nsp14 MTase activity. Alongside these structures, their respective 2D interaction diagrams, derived from docking studies, are displayed. The figure also depicts the putative 3D poses these compounds assume within nsp14’s lateral cavity.

### B. Experimental Evaluation of Lead Compounds

In this study, we harnessed both the precision and accuracy of a state-of-the-art high-throughput radioactivity-based assay [21], to experimentally test the MTase activity of nsp14 under the influence of our selected compounds. Out of the 40 assessed compounds, seven manifested inhibitory properties (Table I). Among these, NSC76988 and NSC620333 emerged as particularly promising candidates, due to their relatively low IC_50_ values 6 and 5 *μ*M, respectively, suggestive of a potent inhibitory effect. To further substantiate these findings and confirm the binding of these inhibitors to nsp14, we employed surface plasmon resonance (SPR). [26]. This technique was leveraged to investigate the binding interactions of the top four compounds from the radioactivity-based assay. Figure 2 provides detailed insight into these molecular interactions. It presents the 2D structural diagrams of the top four experimentally validated compounds that inhibit MTase activity. Alongside these are the 2D interaction diagrams derived from docking studies, and 3D poses showing how these compounds fit within the lateral cavity of nsp14. Interestingly, of these four inhibitors, NSC620333 had the lowest *K*_*D*_ value of 544 ± 22 nM (cf. Figures3 and S1). This observation points to the potential of NSC620333 as an effective nsp14 inhibitor. Moreover, the results from batch re-testing with a purified sample of NSC620333 provide compelling reinforcement of this proposition, as depicted in Supplementary Figures S3 and S6.

**FIG. 3.**
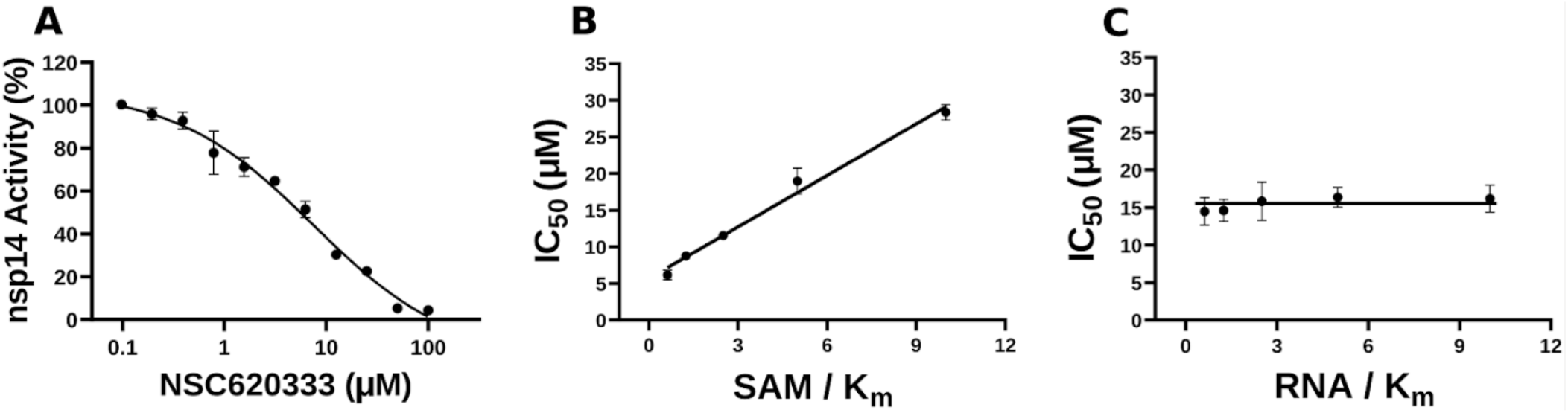
Inhibition of nsp14 MTase activity by NSC620333. **(A)** The IC_50_ value was determined for NSC620333 at *K*_*m*_ of both SAM and RNA substrates (5.3 *±* 1.0 *μM*, Hill Slope: *−*0.9). The mechanism of action (MOA) of NSC620333 was also determined by IC_50_ determination for NSC620333 at **(B)** varying concentrations of SAM (from 0.15 to 2.5 *μM*) at fixed concentration of RNA substrate (0.25 *μM*, 5x *K*_*m*_) and **(C)** varying concentrations of RNA substrate (from 31 to 500 *nM*) at fixed concentration of SAM (1.25 *μM*, 5x *K*_*m*_). All values are presented as mean *±* standard deviation of three independent experiments (*n* = 3). NSC620333 is a SAM competitive nsp14 inhibitor (B), and noncompetitive with respect to RNA substrate (C).

**TABLE I.** High-throughput Screening Results of top 40 Compounds obtained for the Inhibition of nsp14 Methyltransferase (MTase) Activity. This table summarizes the inhibitory effects of the top 40 compounds on nsp14 MTase activity as assessed by high-throughput screening. For each compound, data from two replicate experiments are included, revealing the inhibition percentages at a compound concentration of 100*μ*M. The IC_50_ values, indicating the concentration at which half-maximal inhibition is achieved, are reported for the most promising compounds. The table also provides the observed quenching effects of each compound. The term “ND” indicates that the value is not determined.

| Compound | nsp14 Inhibition % (100μM) | | IC<sub>50</sub> (μM) | Quenching effect % |
| --- | --- | --- | --- | --- |
|  | Exp 1 | Exp 2 |  |  |
| NSC76988 | 100 | 99 | 6 | 8 |
| NSC620333 | 100 | 99 | 5 | 23 |
| NSC77131 | 99 | 99 | 9 | 52 |
| NSC400718 | 89 | 88 | 38 | 23 |
| NSC34443 | 86 | 81 | 57 | -7 |
| NSC630814 | 84 | 82 | ND | 75 |
| NSC102798 | 77 | 74 | ND | 61 |
| NSC4348 | 73 | 61 | 34 | 9 |
| NSC400937 | 73 | 66 | ND | 70 |
| NSC4624 | 65 | 47 | ND | 49 |
| NSC2269 | 64 | 60 | 27 | 1 |
| NSC255523 | 59 | 33 | ND | -7 |
| NSC670682 | 50 | 54 | ND | 43 |
| NSC99790 | 46 | 36 | ND | 33 |
| NSC646375 | 26 | 20 | ND | 1 |
| NSC27605 | 26 | 10 | ND | 2 |
| NSC107661 | 18 | 7 | ND | -2 |
| NSC293892 | 15 | 3 | ND | -5 |
| NSC44037 | 15 | 9 | ND | 8 |
| NSC77680 | 13 | 7 | ND | 21 |
| NSC80136 | 13 | 2 | ND | -3 |
| NSC114010 | 11 | -3 | ND | -2 |
| NSC268226 | 9 | 0 | ND | 3 |
| NSC163444 | 9 | 5 | ND | 2 |
| NSC377438 | 8 | -3 | ND | 17 |
| NSC131119 | 8 | 0 | ND | 14 |
| NSC39302 | 7 | -2 | ND | 12 |
| NSC317609 | 7 | 1 | ND | 8 |
| NSC232469 | 7 | -7 | ND | -4 |
| NSC655184 | 7 | -11 | ND | 5 |
| NSC92432 | 6 | -1 | ND | 8 |
| NSC137050 | 6 | -1 | ND | 3 |
| NSC158437 | 5 | 1 | ND | 15 |
| NSC613624 | 4 | -1 | ND | 6 |
| NSC60360 | 4 | -4 | ND | 10 |
| NSC330685 | 4 | 2 | ND | 1 |
| NSC54251 | 4 | -4 | ND | -1 |
| NSC313453 | 3 | -5 | ND | 5 |
| NSC132916 | -1 | -3 | ND | 3 |

In the quest to further understand the specificity of NSC620333, we evaluated its inhibitory activity against a diverse panel of 33 human RNA-, DNA-, and protein-MTases (Supplementary Table S3). A critical part of our investigation focused on potential off-target effects within the host to ensure the safety and efficacy of the compounds. Our analyses revealed that NSC620333 selectively inhibited only human RNMT [27] (N^7^ guanosine RNA methyltransferase) and PRMT7 [28] (protein arginine methyltransferase 7), with IC_50_ values of 8.6 *±* 1.3 and 7 *±* 1.5 *μ*M, respectively (Supplementary Figure S2). Notably, RNMT is involved in the process of mRNA capping, which is essential for maintaining mRNA stability, facilitating its export, and ensuring efficient translation [29]. In contrast, PRMT7 participates in the methylation of arginine residues on specific proteins, thereby impacting cellular processes such as gene transcription, DNA repair, and signal transduction [30]. Furthermore, to extrapolate our *in vitro* findings to a physiologically-relevant context, we tested NSC620333 and NSC76988 in an in *vivo assay*. This experiment was based on a lung epithelial SARS-CoV-2 infection model using Calu-3 cells. Encouragingly, both compounds demonstrated anti-SARS-CoV-2 activity (Supplementary Figure S5), providing supportive evidence for their potential as therapeutic agents. Taken together, the data from all three complementary assays converges towards a common conclusion: NSC620333 stands out as a potential therapeutic candidate against SARS-CoV-2. This compound exhibited a potent inhibitory effect, significant binding to nsp14, favorable selectivity, and in vivo efficacy. These promising attributes warrant further investigations into its potential for clinical application in SARS-CoV-2 treatment. On a separate note, it is interesting to observe that while no direct binding was detected between NSC76988 and nsp14, the compound displayed notable inhibitory effects. However, the mechanism of such inhibition is unclear.

### C. MD Simulations of nsp14

Structural insights into nsp14 were obtained through molecular dynamics (MD) simulations performed under three different conditions: (i) the protein only, (ii) the protein in complex with NSC620333, and (iii) nsp14 in/out of complex with metal coordination complexes. Our findings are presented as follows:

#### 1. Protein dynamics

Investigations into ligand binding within the MTase lateral cavity were conducted via numerous extended molecular dynamics simulations of the nsp14-nsp10 complex, all performed under a consistent thermal condition of 303.15 *K* (details in Methods: Molecular Dynamics Simulations V G). Throughout these simulations, we monitored the average motion of residues located within a 5-angstrom radius of the SS148 binding region, as depicted in Figure 4(A). A thorough evaluation of the complete motion of the associated residues is presented in Supplementary Figures S7. Interestingly, we found that residues Phe367, Tyr368, and Gln354 on average exhibited the most significant movement across all simulations. These residues displayed considerable stabilization when NSC620333 was present in the binding pocket. Specifically, the root-mean-square deviation (RMSD) movement of Phe367, Tyr368, and Gln354 reduced to 0.25, 0.29, and 0.26 angstroms respectively in the presence of NSC620333. Significantly, we observed a limited number of conformational changes in the nsp10 protein when it was complexed with nsp14 (see Figure 4(B) (Left)), in-dicating a high level of interaction stability between the two proteins based on our sampling. Figure 4(B) (Middle/Right) showcases the key residues involved in this interaction. Specifically, Phe24, Glu11, His85, Thr10, and Val26 were identified as the most crucial residues facilitating this interaction.

**FIG. 4.**
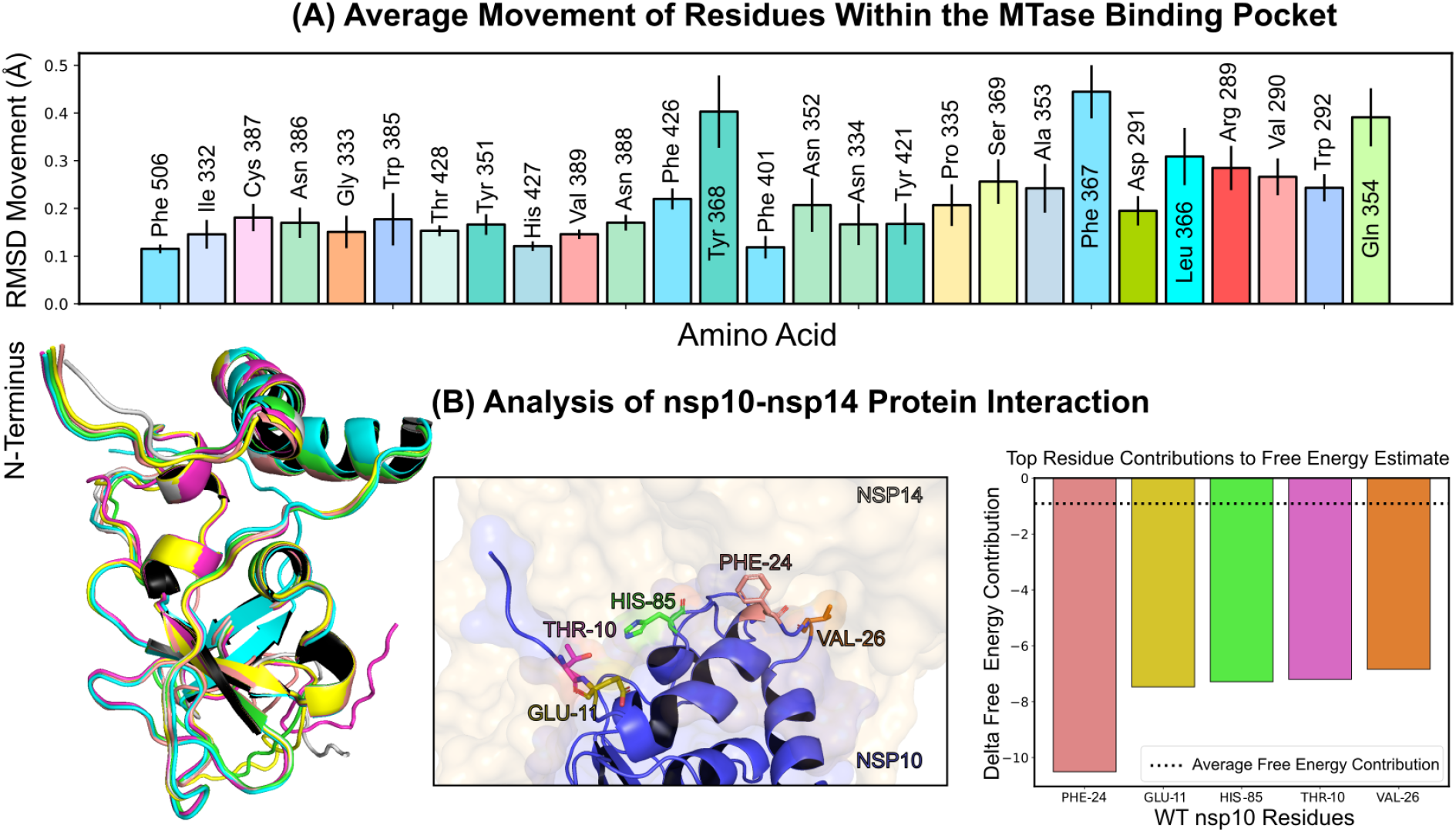
Protein Conformation and Interaction Dynamics from Molecular Simulations. **(A)** Bar charts presenting the mean displacement (expressed in angstroms) of amino acid residues within a 5-angstrom vicinity of the binding pocket, derived from 1-microsecond molecular dynamics simulations conducted at a stable temperature of 303.15 *K*. The error bars indicate the standard deviation of the mean, based on five distinct simulation runs. **(B)** (Left) A graphical representation of the most notable conformational shifts in nsp10, as determined through a series of MD-simulations involving nsp10-14, with only minor changes due to the consistent stability of the nsp10-nsp14 interaction. (Middle) Select residues at the nsp10 binding interface making the most significant contribution to the interaction with nsp16. (Right) Breakdown of individual residues of nsp10 with the highest free energy contributions (kcal/mol) to the overall binding free energy estimation of nsp10-14 interactions; the average contribution from a single residue is indicated by a dotted line.

#### 2. Conformational Shifts

To gain a more comprehensive understanding of the conformational dynamics of the nsp14 complex, we conducted an in-depth analysis of the gathered molecular dynamics simulations [16]. This process involved amalgamating data from receptor-ligand simulations to scrutinize the specific dynamics of the MTase binding cavity and, in a separate analysis, the overall dynamics of the complex backbone across independent trajectories [31]. We employed time-lagged independent component analysis (tICA) to project pertinent trajectory features onto a lower-dimensionality landscape, which facilitated clustering and visualization of the sampled metastable states [32]. The features of the binding pocket were defined by pairwise distances between the heavy atoms of the NSC620333 ligand and the backbone *α*-carbons of 27 crucial residues within the MTase lateral cavity. The analysis of global dynamics similarly employed pairwise distances between *α*-carbons. The generated projections were clustered using the K-means algorithm [33], resulting in eight states that spanned the sampled landscape and were aligned for comparative visualization. In both instances, the cluster with the highest population (comprising approximately 60% of the frames) closely mirrored the anticipated native state, with one or two additional states constituting the majority of the remaining configurations. The visualization of the states underpinning global dynamics revealed that nsp10 and the exoribonuclease domain of nsp14 remained stable. In contrast, the MTase domain exhibited a bent configuration in a minor proportion of frames. The comparative analysis between trajectories both devoid of metal ions and those inclusive of metal ions yielded consistent overall dynamics (see Methods: Molecular Dynamics Simulations V G). However, a larger bend of the main alpha-helix MTase was observable in a more significant population for ion-inclusive trajectories, as evidenced by Cluster 5 and Cluster 7 in Figure 5 (4.6% vs. approximately 1%). This analysis showcases the influence of metal ion presence on the conformational flexibility of the MTase domain, shedding light on the critical role of coordination complex ions in regulating protein structure and dynamics.

**FIG. 5.**
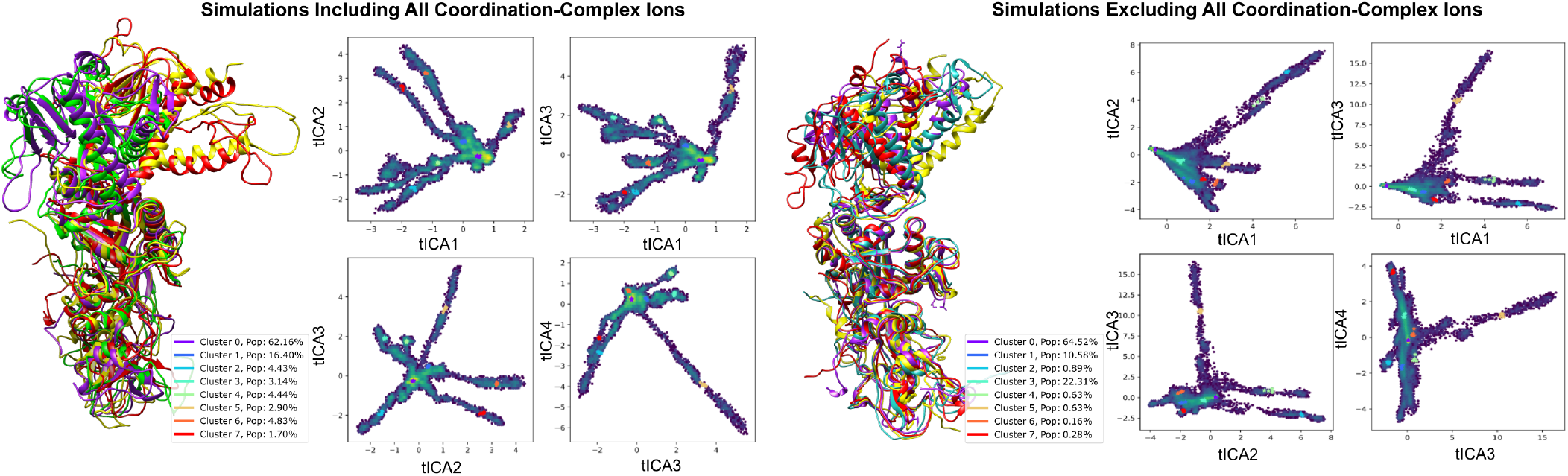
Analyzing Nsp14-nsp10 Conformational Variability through tICA Projections Influenced by Coordination Complexes. This figure delineates the conformational differences in the nsp10-nsp14 complex under ion-devoid and ion-inclusive states, emphasizing structures from four representative cluster centers with high variance. Trajectories were derived from independent runs between 200 ns *−* 1 *μ*s, focusing on pairwise distances between alternating backbone *α*-carbons to determine simulation features and enhance the sampling of diverse conformations.

#### 3. Structural Insights into Inhibition

The application of tICA decomposition on protein-ligand interactions in the simulations identified a variety of binding poses for NSC620333 within the nsp14 MTase binding cavity. From this analysis, eight distinct clusters emerged, which were further refined into three primary states, each visualized in Figure 6. Detailed examination of these states facilitated the identification of specific interactions that contribute to the increased binding affinity of NSC620333, as supported by Wagner et al. [34]. Of particular interest were the hydrogen bonds observed to form from ASN538 to the purine group of the ligand, and the favorable lipophilic interactions of the chlorobenzene component of the ligand with the lipophilic subpocket, predominantly in Cluster 2 and Cluster 6 [35]. These interactions seem to play a crucial role in establishing a more stable binding configuration. Moreover, the varied poses of NSC620333 within the binding pocket further underscore the flexibility of the nsp14 MTase binding cavity, a factor that could be key in the design of effective therapeutic agents [36]. The presence of different binding modes, each with unique interaction profiles, may influence the ligand’s functional impact on the protein, potentially offering multiple pathways for pharmacological intervention [37].

**FIG. 6.**
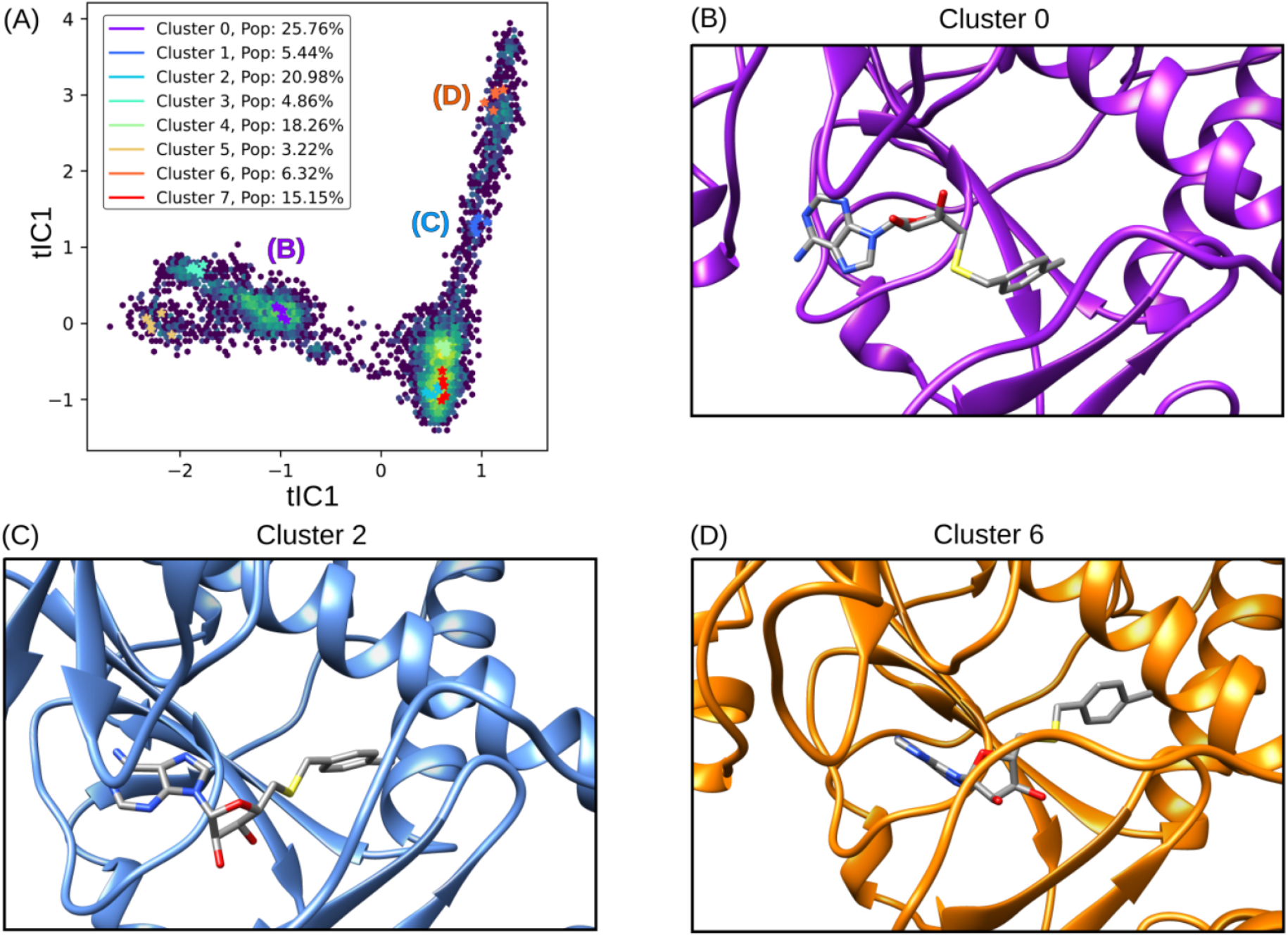
tICA Decomposition and Representative Structures of NSC620333 Alternative Binding Poses. The figure illustrates the tICA decomposition and associated structures of various binding poses of NSC620333, derived from a combination of trajectories that incorporate unbiased molecular dynamics simulations. The trajectories were featurized by calculating pairwise distances between heavy ligand atoms and backbone *α*-carbons of key residues surrounding the binding site. Cluster 2 illustrates the prevalent metastable binding mode observed, while Cluster 0 and Cluster 6 depict twisted and inward binding modes, respectively.

### III. CRYSTAL STRUCTURE OF NSP14 MTASE IN COMPLEX WITH SS148

To uncover the atomic details of the nsp14-SS148 interaction, we performed the crystallographic analysis of the nsp14/SS148 complex. We employed the fusion protein-assisted crystallization approach using the MTase domain of nsp14 (residues 300-527) N-terminally fused to a small crystallization tag TELSAM developed by Kottur et al. [38]. The nsp14 MTase-TELSAM/SS148 crystals belonged to the hexagonal P65 space group and diffracted to the 2.6 Å resolution. The structure of the nsp14 MTase-TELSAM/SS148 complex was subsequently solved by molecular replacement and further refined to good R factors and geometry, as summarized in Supplementary Table S4.

The SS148 ligand was bound to the SAM/SAH-binding site of nsp14 as expected (Fig 7a). The structure of SS148 is derived from SAH and thus, the mechanism of its interaction with nsp14 is also similar to SAH. The nsp14-SS148 interaction is mediated by multiple interactions including both direct (via Asp352, Ala353, Tyr368, and Trp385) and indirect water-mediated hydrogen bonds (via Gln313, Asp331, and Ile332) (Fig 7b). The amino acid moiety of SS148 binds to nsp14 amino acid residues Gln313, Trp385, and Asp331, while the central ribose-derived moiety of nsp14 interacts mainly with Asp352, and the basic adenine-derived moiety binds to Ala353 and Tyr368.

**FIG. 7.**
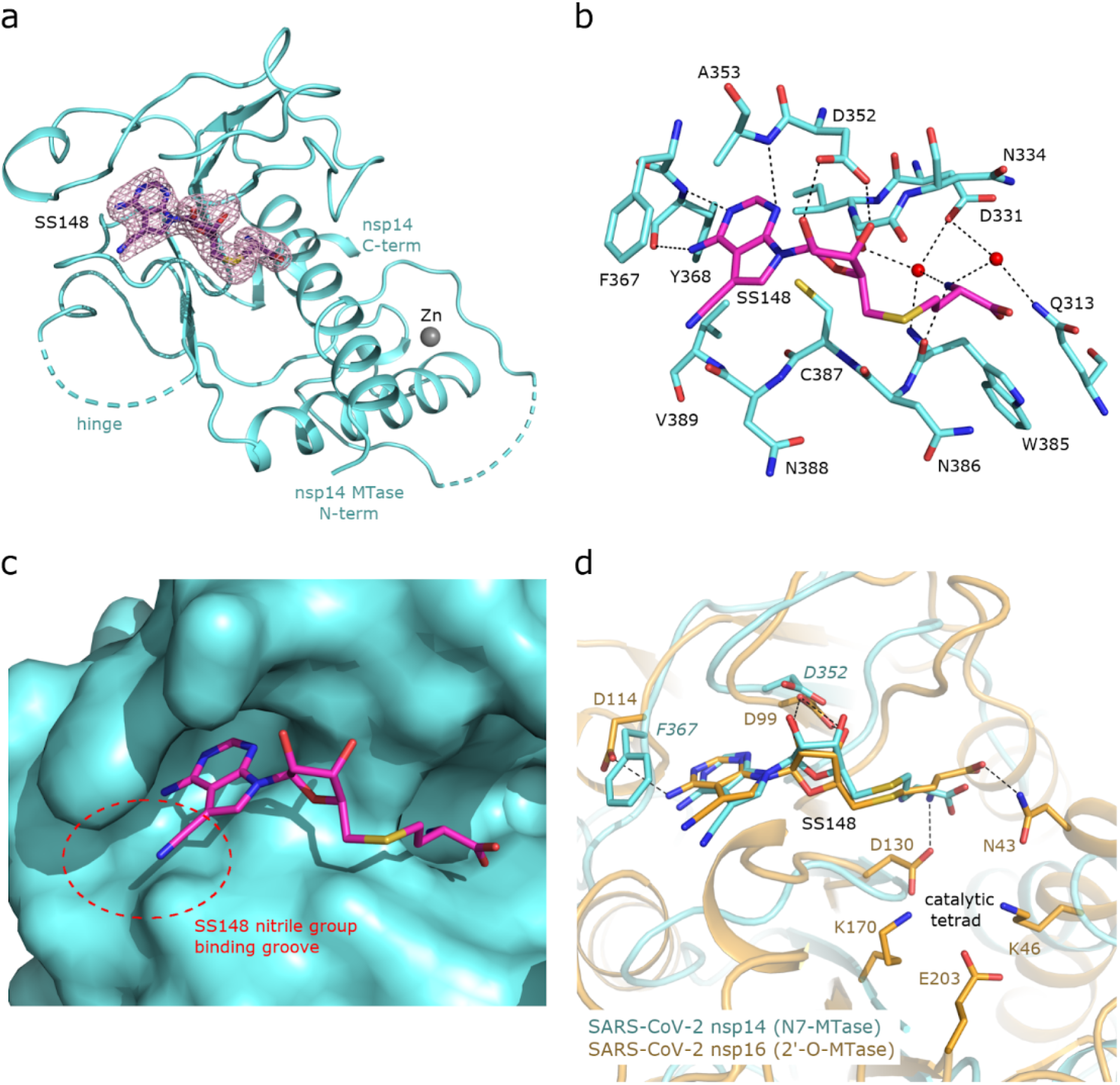
Crystal structure of nsp14 MTase in complex with SS148. **(a)**, Overall view of the nsp14/SS148 complex. The protein backbone of the nsp14 MTase domain is shown in cartoon representation and colored in light blue. The SS148 ligand is shown in stick representation and colored according to elements: carbon, magenta; nitrogen, blue; oxygen, red; sulfur, yellow. The unbiased Fo-Fc omit map contoured at 2s is shown around the SS148 ligand. The TELSAM crystallization tag N-terminally fused to the nsp14 MTase domain is not shown. **(b)**, Detailed view of the SS148 ligand binding site. The SS148 ligand and side chains of selected nsp14 amino acid residues are shown in stick representation, with carbon atoms colored according to the protein assignment and other elements colored as in (a). Water molecules are presented as red spheres; hydrogen atoms are not shown. Selected hydrogen bonds involved in the nsp14–SS148 interaction are depicted as dashed black lines. **(c)**, SS148 ligand binding site with nsp14 shown in surface representation. The nitrile group binding groove on the surface of nsp14 potentially accepting larger substituents is highlighted with a red dashed circle. **(d)**, Structural alignment of the SS148 binding sites of SARS-CoV-2 N7-MTase nsp14 and 2’-O-MTase nsp16. Protein backbones are shown in cartoon representation, while SS148 and side chains of selected residues are shown in stick representation. The carbon atoms of the nsp14/SS148 and nsp16/SS148 complexes are depicted in light blue and yellow, respectively; other elements are colored as in (a).

Compared to SAH, SS148 contains a nitrile group bound to the C7-position of the 7-deaza-adenine heterocyclic moiety. This nitrile group is bound to nsp14 in the groove formed by the nsp14 residues Phe367, Asn388, and Val389 (Fig 7c). Our study documents that introduction of substituents at the C7-position of this 7-deaza-adenine moiety, physically larger than the presented nitrile group, can be a promising strategy potentially leading to development of more potent and more specific nsp14 inhibitors.

SS148 is an inhibitor with dual activity against both SARS-CoV-2 N7-MTase nsp14 and 2’-O-MTase nsp16. Therefore, we compared the SS148 binding site of nsp14 with the previously crystallized SS148 binding site of nsp16 [22] (Fig 7d). Despite significant differences in the organization of the SS148 binding pockets in nsp14 and nsp16, we observed similar conformation of SS148. The conserved catalytic tetrad characteristic for 2’-O-MTases formed by Lys46, Asp130, Lys170, and Glu203 in nsp16 is not present in nsp14.

To judge the reliability of our computational pipeline, we performed re-docking of SS148 onto the nsp14 binding site. Our methodological approach was successful in accurately replicating the binding pose of the SS148 ligand within the nsp14 binding site, showcasing a striking alignment (RMSD 1.426Å) with the experimentally determined crystal structure (depicted in Supplementary Figure S4). The high degree of concordance between the computationally determined re-docking pose and the experimental crystal structure attests to the robustness of our docking pipeline methodology. The agreement between our computational predictions and experimental findings underscores the proficiency of our integrated approach and further substantiates the potential of SS148 as a promising dual inhibitor of SARS-CoV-2 MTases.

## IV. CONCLUSION AND OUTLOOK

Our comprehensive study has provided valuable insights into the complex dynamics of the SARS-CoV-2 nsp14 protein and the therapeutic potential of compounds aimed at inhibiting it. In particular, we found that the compound NSC620333 demonstrates promising attributes as a therapeutic agent against SARS-CoV-2, primarily due to its potent inhibitory effects on the nsp14 Methyltransferase (MTase) activity and its substantial binding affinity to nsp14. The observed in-vivo efficacy and selectivity of this compound for development of therapeutics in treating COVID-19. In our study, we identified several critical residues within the MTase binding pocket of nsp14, and elucidated their dynamic behavior in the presence of potential inhibitors, such as NSC620333. Our findings lay a solid foundation for future drug design efforts, paving the way for the development of novel strategies to disrupt the nsp14 function. This could potentially lead to new therapeutic options for COVID-19.

Interestingly, we observed conformational changes in nsp14, particularly in the presence and absence of metal coordination complexes, which emphasizes the role of this metal ion in the structural dynamics of nsp14. The diverse binding poses of the NSC620333 ligand identified in our study highlight the necessity for a dynamic view of protein-ligand interactions. It suggests that a single static view of a protein-ligand complex is insufficient to capture the full complexity of these interactions, further emphasizing the value of molecular dynamics simulations in comprehending these systems. Our structural studies also revealed the binding mechanism of another potential inhibitor, SS148. By visualizing its binding mode and interactions with nsp14, we proposed that introducing substituents at the C7-position of the 7-deaza-adenine moiety could enhance the potency and specificity of nsp14 inhibitors. The crystal structure also revealed the differences and similarities in the SS148 binding site between nsp14 and nsp16, providing valuable structural insights for drug design.

While our results are promising, they represent only the initial steps in a much longer journey. Further studies are needed to fully understand the therapeutic potential of these compounds. Future work should focus on optimizing these lead compounds, assessing their pharmacokinetic and pharmacodynamic properties, and evaluating their safety and efficacy in pre-clinical and clinical trials. Furthermore, the mechanistic exploration of NSC76988 and its potential allosteric mode of inhibition should also be pursued. Overall, this study underscores the potential of computational methods in understanding the complexities of biological systems and guiding the development of new therapeutics. We hope that our findings will inspire and inform future research efforts towards the development of effective treatments for COVID-19 and potentially other related viral threats.

## V. METHODS

### A. Docking

The Cryo-EM structure of SARS-CoV-2 nsp10-nsp14 (PDB:7N0B [39]) was utilized as a reference for molecular docking. The receptor was prepared through Schrödinger’s protein preparation tool [40], which involved a series of critical steps. These steps included modeling missing residues, capping missing termini, removing water molecules, and assigning protonation states. The ligand binding site was identified by superimposing SAM/SAH (PDB: 5C8S/5C8T [41]) onto the Cryo-EM structure. Upon completion of the overlay, a minimization was performed using the OPLS4 force field [42].

The process aimed to optimize the overall structure of the receptor-ligand complex, minimize potential clashes, and maximize binding interactions. For the molecular docking process, we employed Glide-SP and Glide-XP software [25], integrated within the VirtualFlow Unity [20] (VFU) pipeline (https://github.com/VirtualFlow/VFU). In the interest of scientific inclusivity, we provide a VFU setup that utilizes QuickVina [43] and Smina [44], two open-source docking programs.

The NCI Open Compound collection was procured from NCBI PubChem (https://pubchem.ncbi.nlm.nih.gov/) by selecting “DTP/NCI” as the Data Source. The provided isomeric smiles were employed as starting points for processing the initial 250,000 molecules. At the stage of identifying the top-100 molecules, the molecules were manually inspected to ensure correct stereochemistry.

Diversity for a set of molecules was calculated as:

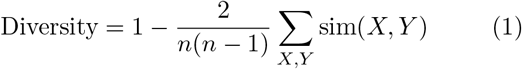

The expression sim(*X, Y*) computes the pairwise molecular similarity for all *n* structures calculated as the Tanimoto distance of the Morgan fingerprint (calculated with a radius of size 3 and a 2048 bit size) [45].

### B. MM/GBSA Calculation

The AMBER ff14SB force field [46] and the General AMBER force field (GAFF2) [47] were employed for parameterizing proteins and ligands, respectively, in the complex systems. The antechamber module [48] was utilized to calculate the partial atomic charges using the AM1-BCC charge model for ligand molecules. To eliminate bad steric contacts, the systems were subjected to energy minimization with the steepest descent algorithm and conjugate gradient methods, without restraints. This process involved 2,500 iterations using the Sander MD engine [49]. The MM/GBSA energies were evaluated using the MMPBSA.py [14] script, which is part of the AmberTools21 package [50].

### C. Protein Expression and Purification

The SARS-CoV-2 nsp14 MTase domain-encoding sequence (GenBank: YP 009725309.1, residues 300-527) was codon-optimized for expression in *E. coli* and synthesized by GeneArt (Thermo Scientific). The gene was cloned into a modified pRSFDuet vector containing an N-terminal hexahistidine (His6) purification tag, followed by a SUMO solubility and folding tag, and a TELSAM crystallization tag [38]. The plasmid was transformed into *E. coli* BL21 DE3 NiCo bacterial strain (New England Biolabs), and the protein was overexpressed using autoinduction ZY medium. Cells were collected by centrifugation and resuspended in lysis buffer (50 mM Tris, pH 8.0, 400 mM NaCl, 20 mM imidazole, 10 mM MgCl_2_, 10 *μ*M ZnCl_2_, 3 mM *β*-mercaptoethanol, and 250U of DNA endonuclease DENERASE (c-LEcta)). The cells were then sonicated using the Q700 Sonicator instrument (QSonica). The lysate was subsequently cleared by centrifugation, and the supernatant was incubated with Ni-NTA agarose (Thermo Scientific), followed by extensive washing with lysis buffer. The protein was finally eluted using lysis buffer supplemented with 300 mM imidazole. Post-elution, the protein was treated with Ulp1 protease to cleave off the His6-SUMO tag. The nsp14 MTase-TELSAM protein was further purified by size exclusion chromatography using Superdex 200 16/600 (GE Healthcare) pre-equilibrated with size-exclusion buffer (25 mM Tris pH 8.3, 200 mM KCl, and 2 mM TCEP). Fractions containing the nsp14 MTase-TELSAM protein were concentrated to 4 mg/ml, flash-frozen, and stored at *−*80^*°C*^C.

### D. Crystallization and Crystallographic Analysis

Crystallization trials were conducted with a concentration of 4 mg/ml of the nsp14 MTase-TELSAM protein, supplemented with a twofold molar excess of the SS148 ligand. Crystals were observed to grow within two days at 18^*°C*^C. This was achieved in a sitting drop composed of a 200 nl mixture of the protein and 200 nl of the mother liquor, which contained 100 mM bicine/Trizma pH 8.5; 10% w/v PEG 20,000; 20% v/v PEG MME 550; 20 mM D-glucose; 20 mM D-mannose; 20 mM D-galactose; 20 mM L-fucose; 20 mM D-xylose; 20 mM N-acetyl-D-glucosamine.

The crystallographic dataset was collected from a single crystal at the BL14.1 beamline, located at the BESSY II electron storage ring operated by Helmholtz-Zentrum Berlin [51]. Data integration and scaling were performed using XDS [52]. Molecular replacement was used to solve the structure of the nsp14 MTase-TELSAM/SS148 complex, with the nsp14 MTase-TELSAM/sinefungin complex structure serving as the search model (PDB entry 7TW9 [38]). Phaser from the Phenix package was used for the initial model [53, 54]. Model improvement was accomplished via automatic model refinement with Phenix.refine [55] and manual model building with Coot [56]. Statistics regarding data collection, structure solution, and refinement are provided in Supplementary Table S4. Structural figures were generated using PyMOL Molecular Graphics System v2.5 (Schrödinger, LLC) [57]. The atomic coordinates and structural factors have been deposited in the Protein Data Bank (https://www.rcsb.org) under the accession code 8BWU.

### E. Radioactivity-based Assay Conditions

The assay conditions for nsp14 were as follows: The final concentrations in the reaction mixture were 1.5 nM for nsp14, 50 nM for Biotin-RNA (*5’GpppACCCCCCCCC-Biotin 3’*), and 250 nM for ^3^H-SAM. The incubation was carried out for 20 minutes at 23°C. The buffer used contained 20 mM Tris HCl at pH 7.5, 250 *μ*M MgCl_2_, 5 mM DTT, and 0.01% Triton X-100, with an additional 5% DMSO.

### F. SPR Conditions

For Surface Plasmon Resonance (SPR), the full-length biotinylated nsp14 was immobilized on the flow cell of a SA sensor chip in 1x HBS-EP buffer, achieving a response of 8000 RU. The assay was performed using the same buffer but with 2% DMSO added. Single-cycle kinetics were employed with a contact time of 60 seconds and a dissociation time of 120 seconds, at a flow rate of 75 *μ*L/min. NSC620333 was tested at a concentration of 10 *μ*M as the highest concentration, and a dilution factor of 0.25 was used to yield five different concentrations.

### G. Molecular Dynamics Simulations

The crystal structure of nsp14 in complex with nsp10 was obtained from the Protein Data Bank (PDB) under the entry code 7N0B. All missing residues and variants relative to the wild-type were modeled using Schrodinger’s protein preparation tool. The RNA complex was removed for the purposes of our simulations. The system was solvated in a periodic cubic box containing TIP3P water with 150 mM KCl using CHARMM-GUI [58]. The CHARMM36m force field [59] with hydrogen mass repartitioning was employed, and input files were sourced directly from CHARMM-GUI. Molecular dynamics simulations were performed using the Gromacs 2020.3 software package [60]. The simulations employed a Langevin thermostat and a Nosé–Hoover Langevin piston barostat to maintain a pressure of 1 atm. Long-range interactions were treated using the particle mesh Ewald method, and non-bonded interactions were smoothed between 10 to 12 Å. The system underwent energy minimization for 1,500 steps, followed by equilibration to 303.15K with a timestep of 2 fs for 10 ns using the SHAKE and SETTLE algorithms [61]. Production simulations were conducted with a 4 fs timestep for a total of 1.0 *μ*s, with five independent replicates. Simulations that included NSC620333 were parameterized using the Charmm General Force Field (CGenFF) framework [62]. Throughout the simulations, the root-mean-square deviation (RMSD) of the protein residues was calculated using the MDTraj package [63]. All our simulations can be downloaded from: https://drive.google.com/drive/folders/1tKiIsDxBbjZLCxceSoDXlJAczd4UgHIJ?usp=sharing.

### H. Anti-SARS-CoV-2 and Cytotoxicity Assays in Calu-3 Cells

Anti-SARS-CoV-2 activity of NSC620333 and NSC76988 compounds were determined in the human lung adenocarcinoma cell line Calu-3 (ATCC, cat. no. HTB-55) using SARS-CoV-2 (isolate hCoV-19/Czech Repub-lic/NRL 6632 2/2020). Briefly, two-fold eight serial dilutions of the compounds starting from 100 *μM* were added to 40, 000 Calu-3 cells seeded for 1 day in DMEM supplemented with 2% fetal bovine serum, 100 *U* of penicillin/mL and 100 *μg* of streptomycin/mL. After 1 *h* incubation at 37 °C, SARS-CoV-2 was added at a multiplicity of infection (MOI) of 0.03 IU/ml, incubated 3 days at 37 °C, and the virus-induced cytopathic effect (CPE) was quantified by formazan-based cell viability assay (XTT assay). XTT solution was added and after 4 *h* incubation, the absorbance at 450 nm was measured using the EnVision plate reader. After plotting the percentage cell viability versus log_10_ concentrations, the compound concentrations required to reduce viral cytopathic effect by 50% (EC50) were calculated by nonlinear regression. To determine compound cytotoxicity, Calu-3 cells were exposed to the same compound dilutions as above but without virus, and cell viability was determined using XTT assay. The compound concentrations causing 50% reduction in cell viability (CC50) were calculated similarly. Three-fold serial dilutions of Remdesivir from 10 *μM* served as a control in both experiments.

### I. Assay Technique

The MTase activity of nsp14 was measured using a radiometric assay as described previously [21]. Chlorobenzothiophene 3 was tested at various concentrations ranging from 200 nM to 200 *μ*M to determine the halfmaximal inhibitory concentration (IC_50_) value. The final reaction mixture consisted of 1.5 nM nsp14, 250 nM ^3^H-SAM, and 50 nM RNA in buffer (20 mM Tris HCl, pH 7.5, 250 *μ*M MgCl_2_, 5 mM DTT, and 0.01% Triton X-100). The reaction time was 20 min at 23^*°C*^C. Data were fitted to the four-parameter logistic equation using GraphPad Prism 8.

### J. Synthesis of SS148

The synthesis of SS148 (also referred to as CN-SAH in the literature) was carried out according to previously published procedures [64]. Comprehensive synthetic procedures and associated 1H and 13C -NMR data for SS148 can be found in the referenced literature.

## ACKNOWLEDGEMENTS

AK.N. acknowledges funding from the Bio-X Stanford Interdisciplinary Graduate Fellowship (SGIF). Special thanks are extended to A.K. and Michael Bassik for their invaluable assistance in securing the SGIF funding. R.P. acknowledges funding through a Postdoc.Mobility fellowship by the Swiss National Science Foundation (SNSF, Project No. 191127). A.A.-G. thanks Dr. Anders G. Frøseth for his generous support. A.A.-G. also acknowledges the generous support of Natural Resources Canada and the Canada 150 Research Chairs program. M.F.D.H and V.A.V were supported in part by the National Institutes of Health grant 1R01GM123296. One part of the parameterization of simulations was performed on HPC resources supported in part by the National Science Foundation through major research instrumentation grant number 1625061 and by the US Army Research Laboratory under contract number W911NF-16-2-0189. The remaining parameterization was performed on the CB2RR cluster made possible through NIH Research Resource computer instrumentation grant S10-OD020095. S.S. is a Damon Runyon Quantitative Biology Fellow supported by the Damon Runyon Cancer Research Foundation (DRQ-14-22). This research was enabled in part by support provided by Calcul Québec (https://www.calculquebec.ca) and Compute Canada (www.computecanada.ca). HN acknowledges NVIDIA and Intel for funding. MD simulations for all compounds were performed on the Béluga supercomputer situated at the École de technologie supérieure in Montreal. We thank the Helmholtz-Zentrum Berlin für Materialien und Energie for the allocation of synchrotron radiation beamtime. This work was supported by the project National Institute Virology and Bacteriology (Programme EXCELES, Project No. LX22NPO5103) - Funded by the European Union - Next Generation EU. The research was also, in part, made possible thanks to funding provided to the University of Toronto’s Acceleration Consortium by the Canada First Research Excellence Fund (CFREF-2022-00042).

## Supplementary Information

**FIG. S1.**
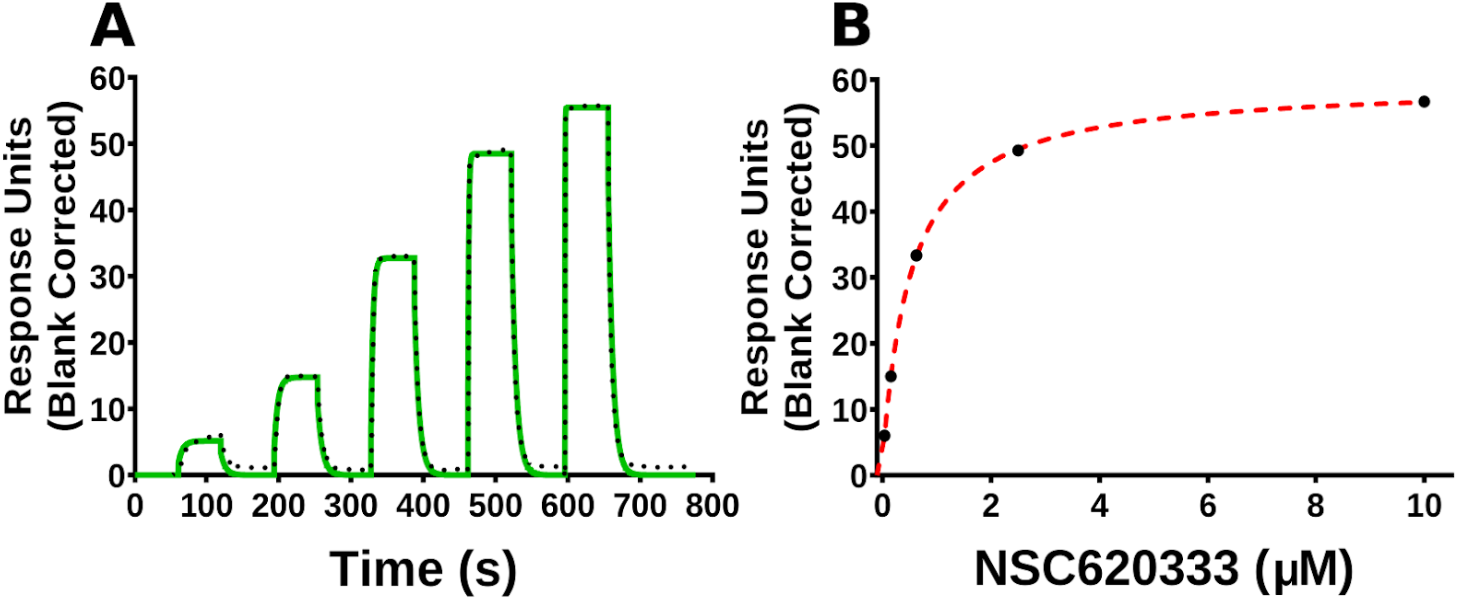
Orthogonal confirmation of NSC620333 binding to nsp14 by SPR. **(A)** A representative sensorgram (solid green) is shown with the kinetic fit (black dots). From kinetic fitting, a *K*_*D*_ value of 427*±*84 *nM, k*_on_ of 3.2*±*0.15*×*10^5^ *M*^−1^ *S*^−1^ and *k*_off_ of 1.3 *±* 0.2 *×* 10^−1^ *S*^−1^ were determined. **(B)** The steady-state response (black circles) obtained from (A) is shown with the steady-state 1:1 binding model fitting (red dashed line). A steady-state *K*_*D*_ value of 544 *±* 22 *nM* (*n* = 3) was also calculated. All values are presented as mean *±* standard deviation of three independent experiments (*n* = 3).

**FIG. S2.**
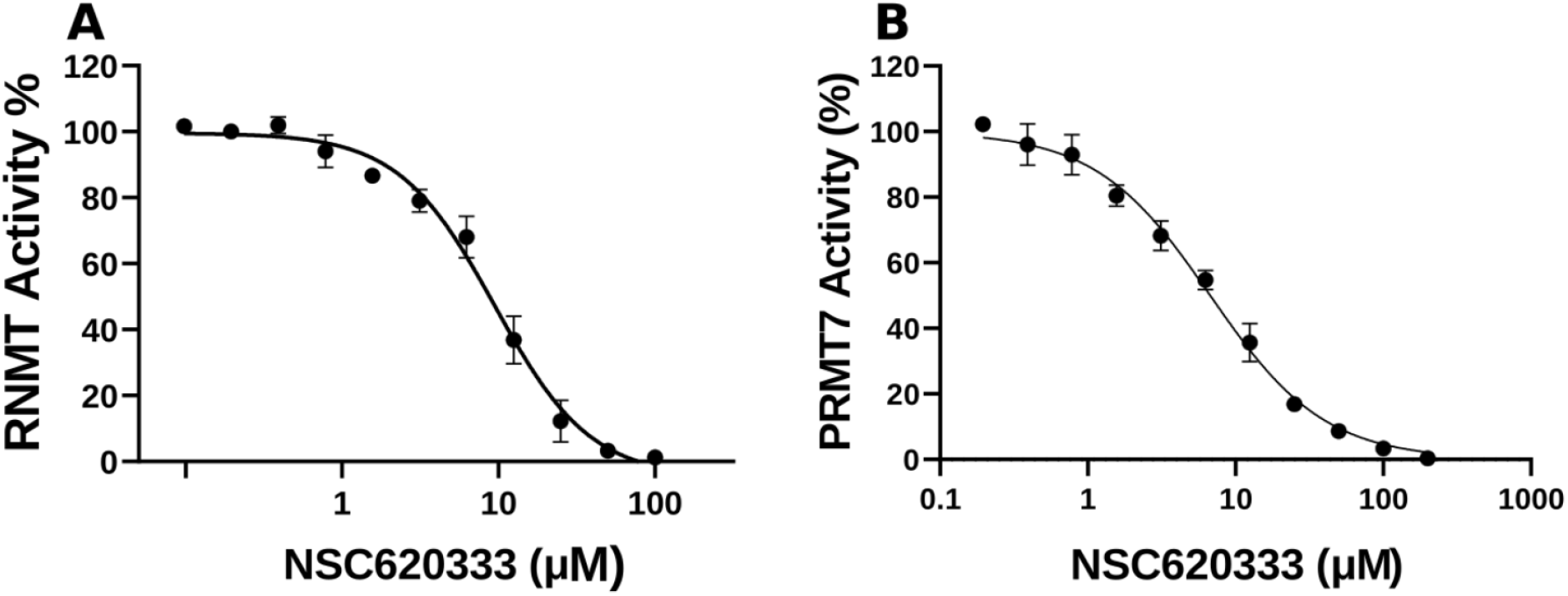
Inhibition of RNMT and PRMT7 activity by NSC620333. (A) The IC_50_ value was determined for NSC620333 to be 8.6 ± 1.3 μM, Hill Slope: -1.5. (B) The IC_50_ value was determined for NSC620333 to be 7.0 ± 0.6 μM, Hill Slope: -1.9.

**FIG. S3.**
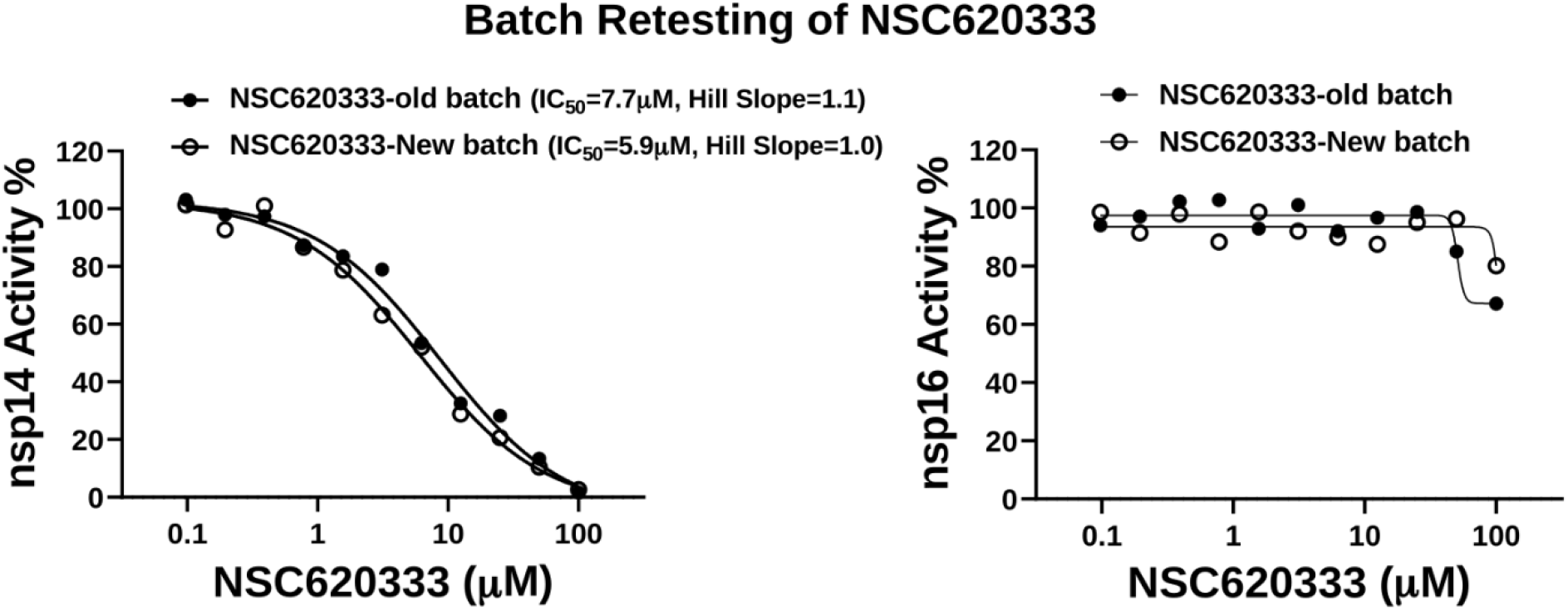
Inhibition Results from Batch Retesting of Purified NSC620333. Outcomes of testing a custom-ordered, highly purified (*>*99%) sample of NSC620333. The exclusion of impurities culminated in a more pronounced inhibition of nsp14, underscoring the potency of the purified compound in the regulation of this complex.

**FIG. S4.**
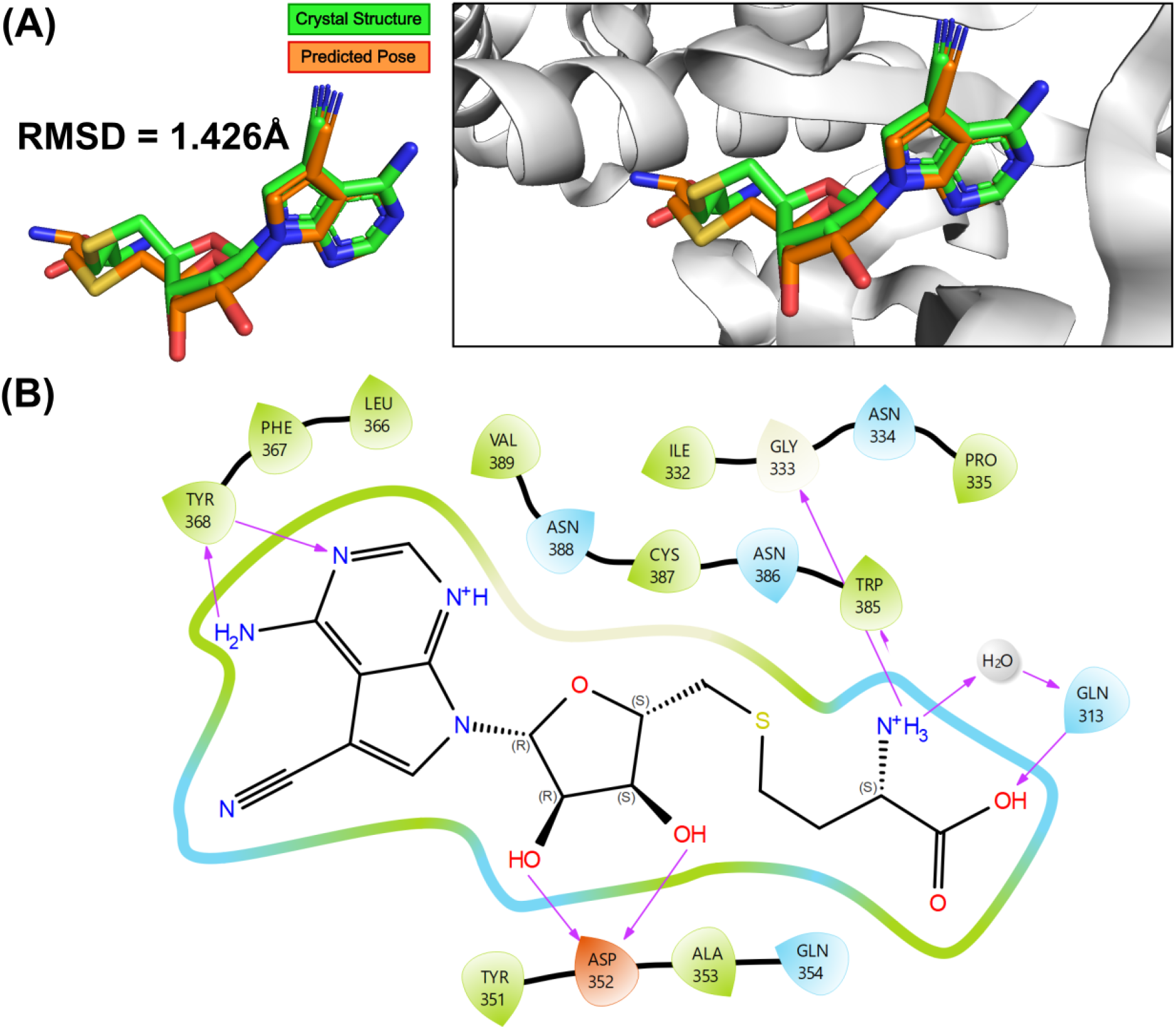
Comparative Analysis of the Crystal Structure and Predicted Docking Pose of SS148 in Complex with nsp14. **(A)** Superimposition of SS148’s actual crystal structure (depicted in green) and its predicted docking pose (depicted in orange), highlighting the high overlap between prediction and experiment. A Root Mean Square Deviation (RMSD) of 1.426Å is observed. **(B)** Two-Dimensional Interaction Diagram depicting the complex formation of SS148 with the nsp14 Methyltransferase (MTase), as per the actual crystal structure.

**FIG. S5.**
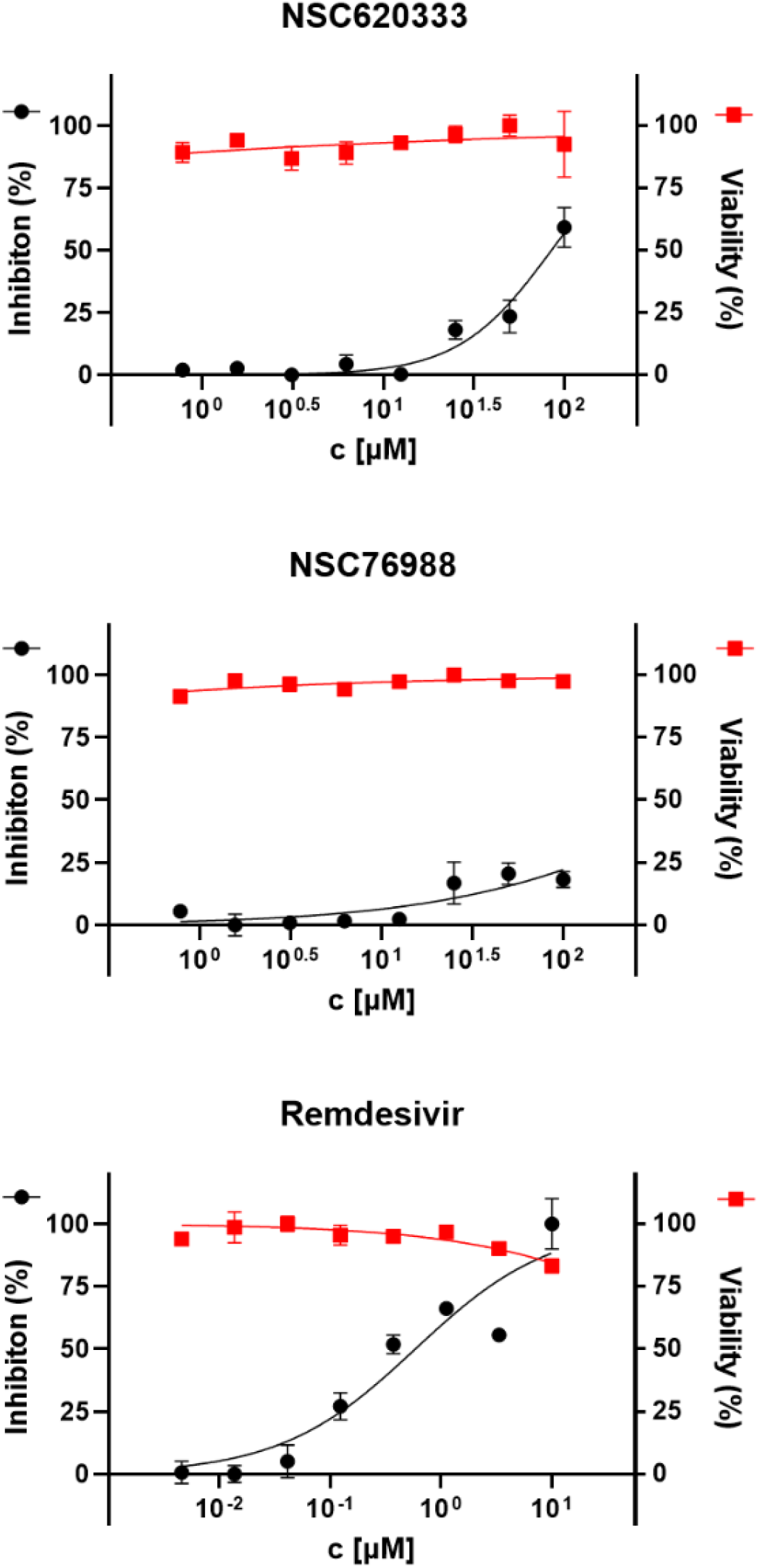
Anti-SARS-CoV-2 activity and cytotoxicity of NSC620333 and NSC76988 in Calu-3 cells. Dose-response curve analysis of anti-SARS-CoV-2 activity (black circle) and cytotoxicity (red square) of NSC620333 and NSC76988 in Calu-3 cells. Remdesivir served as a control. All values are presented as mean ± standard deviations from experiments performed in triplicate.

**FIG. S6.**
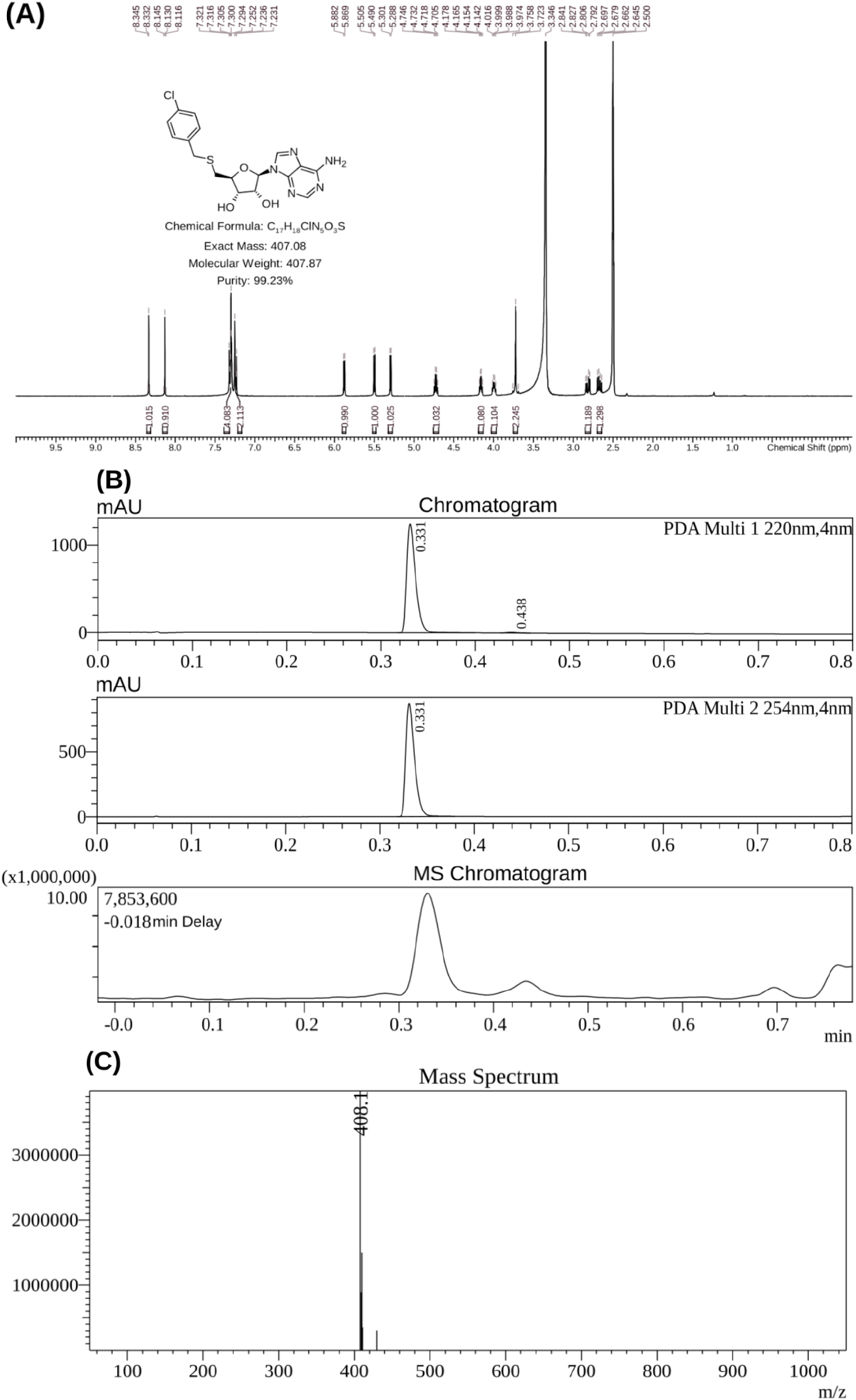
Characterization data for chlorobenzothiophene 3. A. H-NMR spectrum. B. HPLC traces with both UV and MS detectors. C. High-resolution mass spectrum.

**FIG. S7.**
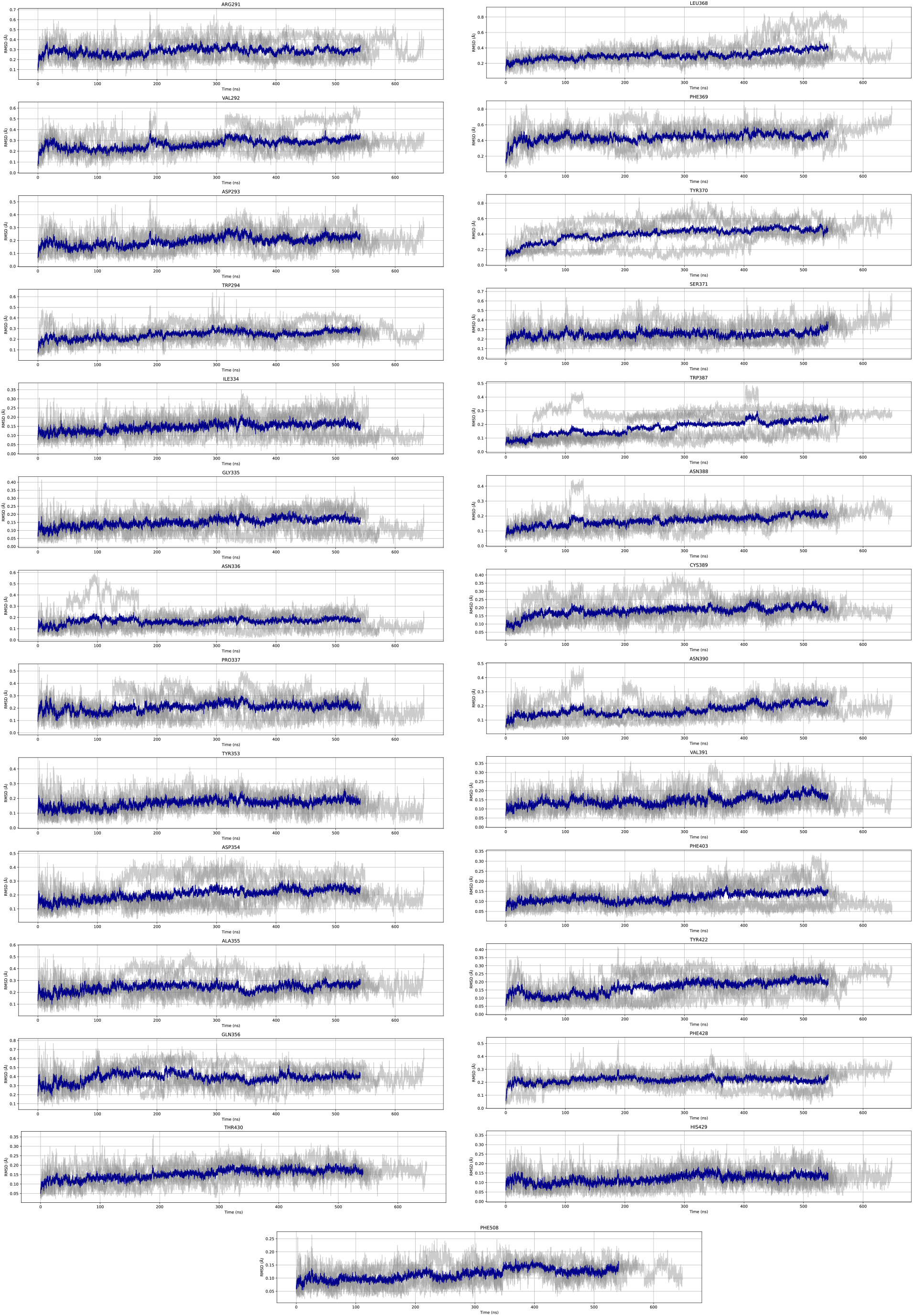
RMSD Variation of Residues in the MTase Lateral Binding Pocket Across 300.15K Simulations. All-atom root mean square deviation (RMSD) movement of residues within the MTase lateral binding pocket during 300.15K simulations across five replicates, relative to the initial conformational structure. The RMSD values are calculated with respect to the initial static structure of the protein-ligand complex, serving as a baseline for measuring conformational changes. The individual RMSD movements for each replicate are depicted as gray lines, illustrating the diversity of conformational changes across the different simulations. The average RMSD movement, plotted in blue, provides a visual summary of the typical movement exhibited by these residues in comparison to the baseline structure.

**TABLE S2.**
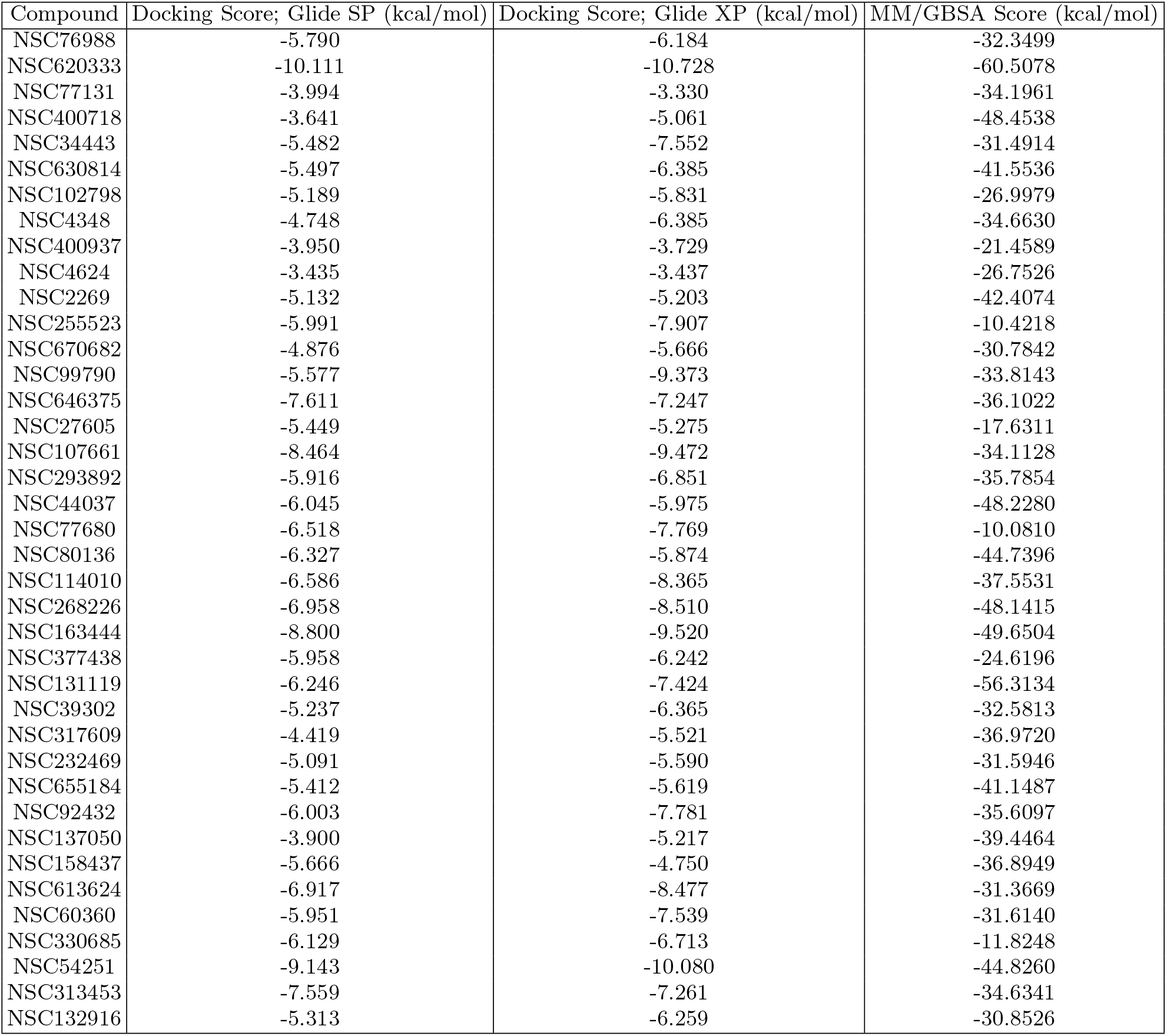
Docking and MM/GBSA Scores of Lead Compounds with nsp14. This figure presents the docking and MM/GBSA scores for 40 lead compounds in complex with nsp14, calculated using three different computational methods. The methods, in increasing order of their predictive capabilities, are Glide SP, Glide XP (both docking-based methods), and MM/GBSA. It should be noted that the MM/GBSA scores are calculated without the entropic term, potentially leading to an overestimation of binding energies. These scores are indicative of the compounds’ potential interactions with nsp14 rather than direct measures of binding affinity.

**TABLE S3.** Inhibitory Activity of NSC620333 on a Diverse Panel of 33 Human Methyltransferases (MTases). Percent inhibitory activity of NSC62033 at a concentration of 10 μM across three replicates for each of the 33 human RNA-, DNA-, and protein-MTases. The average inhibitory activity and standard deviation for each MTase are also provided.

| MTases | Activity (%) at 10 μM | | | |
| --- | --- | --- | --- | --- |
|  | Replicate 1 | Replicate 2 | Replicate 3 | Average |
| G9a | 97 | 79 | 91 | 89 ± 9 |
| GLP | 95 | 105 | 101 | 100 ± 5 |
| SUV39H1 | 99 | 91 | 97 | 96 ± 4 |
| SUV39H2 | 108 | 96 | 106 | 103 ± 7 |
| SUV420H1 | 84 | 97 | 98 | 93 ± 8 |
| SUV420H2 | 96 | 104 | 86 | 96 ± 9 |
| PRMT1 | 97 | 94 | 84 | 91 ± 6 |
| PRMT3 | 75 | 78 | 85 | 79 ± 5 |
| PRMT4 | 96 | 95 | 101 | 97 ± 3 |
| PRMT5 | 69 | 64 | 55 | 63 ± 7 |
| PRMT6 | 97 | 99 | 103 | 100 ± 3 |
| PRMT7 | 35 | 29 | 32 | 32 ± 3 |
| PRMT8 | 97 | 99 | 93 | 96 ± 3 |
| PRMT9 | 118 | 118 | 113 | 117 ± 3 |
| PRDM9 | 110 | 105 | 104 | 106 ± 3 |
| SETDB1 | 104 | 106 | 100 | 103 ± 3 |
| SETD2 | 99 | 103 | 107 | 103 ± 4 |
| SETD7 | 85 | 88 | 87 | 87 ± 1 |
| SETD8 | 79 | 90 | 86 | 85 ± 6 |
| SMYD2 | 104 | 114 | 99 | 105 ± 8 |
| SMYD3 | 101 | 108 | 91 | 100 ± 9 |
| MLL1 | 84 | 87 | 84 | 85 ± 2 |
| MLL3 | 44 | 57 | 53 | 51 ± 6 |
| EZH2 | 101 | 90 | 98 | 96 ± 5 |
| BCDIN3D | 57 | 66 | 64 | 62 ± 5 |
| DOT1L | 103 | 90 | 86 | 93 ± 9 |
| ASH1L | 113 | 97 | 104 | 105 ± 8 |
| NSD1 | 103 | 102 | 107 | 104 ± 2 |
| NSD2 | 100 | 86 | 90 | 92 ± 7 |
| NSD3 | 112 | 104 | 115 | 110 ± 5 |

**TABLE S4.**
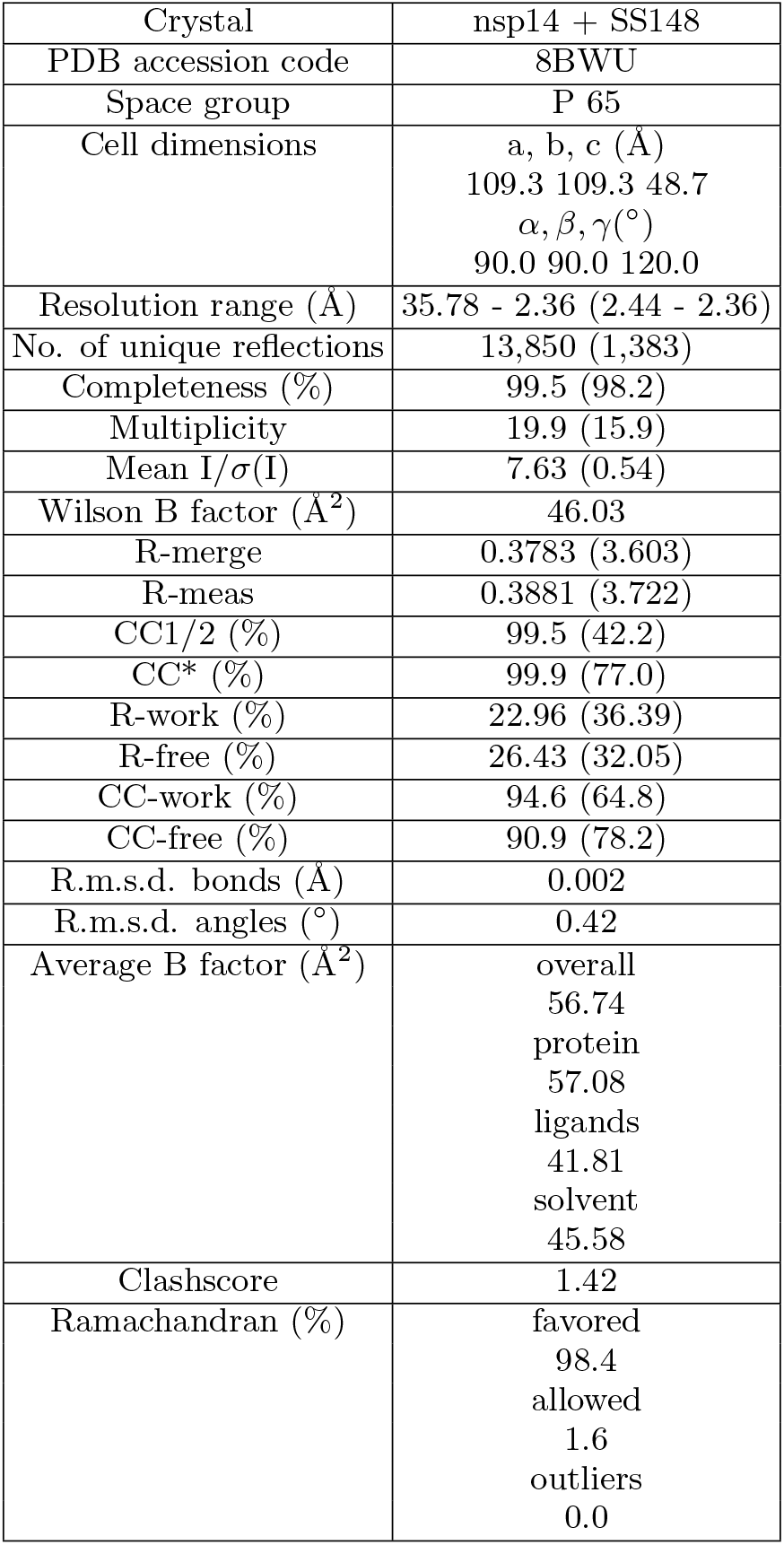
Statistics for data collection and processing, structure solution and refinement of the crystal structure of the nsp14 MTase-TELSAM/SS148 complex. Numbers in parentheses refer to the highest resolution shell. R.m.s.d., root-mean-square deviation.

## References

[1] Catharine I Paules, Hilary D Marston, and Anthony S Fauci. Coronavirus infections—more than just the common cold. Jama, 323(8):707–708, 2020.

[2] World Health Organization et al. The corona virus disease 2019 (covid-19). Brazilian Journal of Implantology and Health Sciences, 4(6):45–46, 2022.

[3] Atul Sharma, Swapnil Tiwari, Manas Kanti Deb, and Jean Louis Marty. Severe acute respiratory syndrome coronavirus-2 (sars-cov-2): a global pandemic and treatment strategies. International journal of antimicrobial agents, 56(2):106054, 2020.

[4] Ahmad Abu Turab Naqvi, Kisa Fatima, Taj Mohammad, Urooj Fatima, Indrakant K Singh, Archana Singh, Shaikh Muhammad Atif, Gururao Hariprasad, Gulam Mustafa Hasan, and Md Imtaiyaz Hassan. Insights into sars-cov-2 genome, structure, evolution, pathogenesis and therapies: Structural genomics approach. Biochimica et Biophysica Acta (BBA)-Molecular Basis of Disease, 1866(10):165878, 2020.

[5] Teresa I Ng, Ivan Correia, Jane Seagal, David A DeGoey, Michael R Schrimpf, David J Hardee, Elizabeth L Noey, and Warren M Kati. Antiviral drug discovery for the treatment of covid-19 infections. Viruses, 14(5):961, 2022.

[6] Mickaël Bouvet, Claire Debarnot, Isabelle Imbert, Barbara Selisko, Eric J Snijder, Bruno Canard, and Etienne Decroly. In vitro reconstitution of sars-coronavirus mrna cap methylation. PLoS pathogens, 6(4):e1000863, 2010.

[7] Yu Chen, Hui Cai, Ji’an Pan, Nian Xiang, Po Tien, Tero Ahola, and Deyin Guo. Functional screen reveals sars coronavirus nonstructural protein nsp14 as a novel cap n7 methyltransferase. Proceedings of the National Academy of Sciences, 106(9):3484–3489, 2009.

[8] Victor M Corman, Doreen Muth, Daniela Niemeyer, and Christian Drosten. Hosts and sources of endemic human coronaviruses. Advances in virus research, 100:163–188, 2018.

[9] Annette von Delft, Matthew D Hall, Ann D Kwong, Lisa A Purcell, Kumar Singh Saikatendu, Uli Schmitz, John A Tallarico, and Alpha A Lee. Accelerating antiviral drug discovery: lessons from covid-19. Nature Reviews Drug Discovery, pages 1–19, 2023.

[10] Wolf-Dietrich Ihlenfeldt, Johannes H Voigt, Bruno Bienfait, Frank Oellien, and Marc C Nicklaus. Enhanced cactvs browser of the open nci database. Journal of chemical information and computer sciences, 42(1):46–57, 2002.

[11] Johannes H Voigt, Bruno Bienfait, Shaomeng Wang, and Marc C Nicklaus. Comparison of the nci open database with seven large chemical structural databases. Journal of chemical information and computer sciences, 41(3):702–712, 2001.

[12] Nataraj S Pagadala, Khajamohiddin Syed, and Jack Tuszynski. Software for molecular docking: a review. Biophysical reviews, 9:91–102, 2017.

[13] AkshatKumar Nigam, Robert Pollice, Gary Tom, Kjell Jorner, Luca A Thiede, Anshul Kundaje, and Alan Aspuru-Guzik. Tartarus: A benchmarking platform for realistic and practical inverse molecular design. arXiv preprint arXiv:2209.12487, 2022.

[14] Bill R Miller III, T Dwight McGee Jr, Jason M Swails, Nadine Homeyer, Holger Gohlke, and Adrian E Roitberg. Mmpbsa. py: an efficient program for end-state free energy calculations. Journal of chemical theory and computation, 8(9):3314–3321, 2012.

[15] Mikko Ylilauri and Olli T Pentik:”ainen. Mmgbsa as a tool to understand the binding affinities of filamin– peptide interactions. Journal of chemical information and modeling, 53(10):2626–2633, 2013.

[16] Scott A Hollingsworth and Ron O Dror. Molecular dynamics simulation for all. Neuron, 99(6):1129–1143, 2018.

[17] J Gelpi. Hospital a, goñi r, orozco m. Molecular dynamics simulations: Advances and applications. Advances and Applications in Bioinformatics and Chemistry, 8:37–47, 2015.

[18] Roni Levin-Konigsberg, Koushambi Mitra, AkshatKumar Nigam, Kaitlyn Spees, Pravin Hivare, Katherine Liu, Anshul Kundaje, Yamuna Krishnan, and Michael Bassik. Slc12a9 is a lysosome-detoxifying ammonium-chloride co-transporter. bioRxiv, pages 2023–05, 2023.

[19] Natacha S Ogando, Jessika C Zevenhoven-Dobbe, Yvonne van der Meer, Peter J Bredenbeek, Clara C Posthuma, and Eric J Snijder. The enzymatic activity of the nsp14 exoribonuclease is critical for replication of mers-cov and sars-cov-2. Journal of virology, 94(23):10–1128, 2020.

[20] Christoph Gorgulla, AkshatKumar Nigam, Matt Koop, Suleyman S Cinaroglu, Christopher Secker, Mohammad Haddadnia, Abhishek Kumar, Yehor Malets, Alexander Hasson, Roni Levin-Konigsberg, et al. Virtualflow 2.0-the next generation drug discovery platform enabling adaptive screens of 69 billion molecules. bioRxiv, pages 2023–04, 2023.

[21] Kanchan Devkota, Matthieu Schapira, Sumera Perveen, Aliakbar Khalili Yazdi, Fengling Li, Irene Chau, Pegah Ghiabi, Taraneh Hajian, Peter Loppnau, Albina Bolotokova, et al. Probing the sam binding site of sarscov-2 nsp14 in vitro using sam competitive inhibitors guides developing selective bisubstrate inhibitors. SLAS DISCOVERY: Advancing the Science of Drug Discovery, 26(9):1200–1211, 2021.

[22] Martin Klima, Aliakbar Khalili Yazdi, Fengling Li, Irene Chau, Taraneh Hajian, Albina Bolotokova, H Ümit Kaniskan, Yulin Han, Ke Wang, Deyao Li, et al. Crystal structure of sars-cov-2 nsp10–nsp16 in complex with small molecule inhibitors, ss148 and wz16. Protein Science, 31(9):e4395, 2022.

[23] Monica Schenone, Vlado Dančík, Bridget K Wagner, and Paul A Clemons. Target identification and mechanism of action in chemical biology and drug discovery. Nature chemical biology, 9(4):232–240, 2013.

[24] Shunzhou Wan, Agastya P Bhati, Stefan J Zasada, and Peter V Coveney. Rapid, accurate, precise and reproducible ligand–protein binding free energy prediction. Interface Focus, 10(6):20200007, 2020.

[25] Richard A Friesner, Jay L Banks, Robert B Murphy, Thomas A Halgren, Jasna J Klicic, Daniel T Mainz, Matthew P Repasky, Eric H Knoll, Mee Shelley, Jason K Perry, et al. Glide: a new approach for rapid, accurate docking and scoring. 1. method and assessment of docking accuracy. Journal of medicinal chemistry, 47(7):1739–1749, 2004.

[26] Jiří Homola and Marek Piliarik. Surface plasmon resonance (SPR) sensors. Springer, 2006.

[27] Dhaval Varshney, Alain-Pierre Petit, Juan A Bueren-Calabuig, Chimed Jansen, Dan A Fletcher, Mark Peggie, Simone Weidlich, Paul Scullion, Andrei V Pisliakov, and Victoria H Cowling. Molecular basis of rna guanine-7 methyltransferase (rnmt) activation by ram. Nucleic acids research, 44(21):10423–10436, 2016.

[28] Kanishk Jain and Steven G Clarke. Prmt7 as a unique member of the protein arginine methyltransferase family: A review. Archives of biochemistry and biophysics, 665:36–45, 2019. 2018.

[29] Annette von Delft, Matthew D Hall, Ann D Kwong, Lisa A Purcell, Kumar Singh Saikatendu, Uli Schmitz, John A Tallarico, and Alpha A Lee. Accelerating antiviral drug discovery: lessons from covid-19. Nature Reviews Drug Discovery, pages 1–19, 2023.

[30] Sharmistha Pal and Saïd Sif. Interplay between chromatin remodelers and protein arginine methyltransferases. Journal of cellular physiology, 213(2):306–315, 2007.

[31] Vijay S Pande, Kyle Beauchamp, and Gregory R Bowman. Everything you wanted to know about markov state models but were afraid to ask. Methods, 52(1):99–105, 2010.

[32] Yusuke Naritomi and Sotaro Fuchigami. Slow dynamics in protein fluctuations revealed by time-structure based independent component analysis: the case of domain motions. The Journal of chemical physics, 134(6), 2011.

[33] IA Hartigan and AS Algoritm. 136: A k-means clustering algorithm/ja hartigan, ma wong. Journal of the Royal Statistical Society Series C (Applied Statistic), 28(1):100–108, 1979.

[34] Jeffrey R Wagner, Christopher T Lee, Jacob D Durrant, Robert D Malmstrom, Victoria A Feher, and Rommie E Amaro. Emerging computational methods for the rational discovery of allosteric drugs. Chemical reviews, 116(11):6370–6390, 2016.

[35] PR Andrews, DJ Craik, and JL Martin. Functional group contributions to drug-receptor interactions. Journal of medicinal chemistry, 27(12):1648–1657, 1984.

[36] Simon J Teague. Implications of protein flexibility for drug discovery. Nature reviews Drug discovery, 2(7):527–541, 2003.

[37] Huameng Li and Chenglong Li. Multiple ligand simultaneous docking: orchestrated dancing of ligands in binding sites of protein. Journal of computational chemistry, 31(10):2014–2022, 2010.

[38] Jithesh Kottur, Olga Rechkoblit, Richard Quintana-Feliciano, Daniela Sciaky, and Aneel K Aggarwal. High-resolution structures of the sars-cov-2 n7-methyltransferase inform therapeutic development. Nature Structural & Molecular Biology, 29(9):850–853, 2022.

[39] Chang Liu, Wei Shi, Scott T Becker, David G Schatz, Bin Liu, and Yang Yang. Structural basis of mismatch recognition by a sars-cov-2 proofreading enzyme. Science, 373(6559):1142–1146, 2021.

[40] Schr odinger Release. 2: Protein preparation wizard, epik, schrödinger, llc, new york, ny, 2021. Impact, Schrödinger, LLC, New York, NY, 2021.

[41] Yuanyuan Ma, Lijie Wu, Neil Shaw, Yan Gao, Jin Wang, Yuna Sun, Zhiyong Lou, Liming Yan, Rongguang Zhang, and Zihe Rao. Structural basis and functional analysis of the sars coronavirus nsp14–nsp10 complex. Proceedings of the National Academy of Sciences, 112(30):9436–9441, 2015.

[42] Chao Lu, Chuanjie Wu, Delaram Ghoreishi, Wei Chen, Lingle Wang, Wolfgang Damm, Gregory A Ross, Markus K Dahlgren, Ellery Russell, Christopher D Von Bargen, et al. Opls4: Improving force field accuracy on challenging regimes of chemical space. Journal of chemical theory and computation, 17(7):4291–4300, 2021.

[43] Amr Alhossary, Stephanus Daniel Handoko, Yuguang Mu, and Chee-Keong Kwoh. Fast, accurate, and reliable molecular docking with quickvina 2. Bioinformatics, 31(13):2214–2216, 2015.

[44] David Ryan Koes, Matthew P Baumgartner, and Carlos J Camacho. Lessons learned in empirical scoring with smina from the csar 2011 benchmarking exercise. Journal of chemical information and modeling, 53(8):1893–1904, 2013.

[45] David Rogers and Mathew Hahn. Extended-connectivity fingerprints. Journal of chemical information and modeling, 50(5):742–754, 2010.

[46] James A Maier, Carmenza Martinez, Koushik Kasavajhala, Lauren Wickstrom, Kevin E Hauser, and Carlos Simmerling. ff14sb: improving the accuracy of protein side chain and backbone parameters from ff99sb. Journal of chemical theory and computation, 11(8):3696–3713, 2015.

[47] Junmei Wang, Romain M Wolf, James W Caldwell, Peter A Kollman, and David A Case. Development and testing of a general amber force field. Journal of computational chemistry, 25(9):1157–1174, 2004.

[48] Junmei Wang, Wei Wang, Peter A Kollman, and David A Case. Antechamber: an accessory software package for molecular mechanical calculations. J. Am. Chem. Soc, 222(1), 2001.

[49] Joseph W Kaus, Levi T Pierce, Ross C Walker, and J Andrew McCammon. Improving the efficiency of free energy calculations in the amber molecular dynamics package. Journal of chemical theory and computation, 9(9):4131–4139, 2013.

[50] David A Case, H Metin Aktulga, Kellon Belfon, Ido Ben-Shalom, Scott R Brozell, David S Cerutti, Thomas E Cheatham III, Vinícius Wilian D Cruzeiro, Tom A Darden, Robert E Duke, et al. Amber 2021. University of California,San Francisco, 2021.

[51] Uwe Mueller, Ronald Förster, Michael Hellmig, Franziska U Huschmann, Alexandra Kastner, Piotr Malecki, Sandra Pühringer, Martin Röwer, Karine Sparta, Michael Steffien, et al. The macromolecular crystallography beamlines at bessy ii of the helmholtzzentrum berlin: Current status and perspectives. The European Physical Journal Plus, 130:1–10, 2015.

[52] Wolfgang Kabsch. xds. Acta Crystallographica Section D: Biological Crystallography, 66(2):125–132, 2010.

[53] Airlie J McCoy, Ralf W Grosse-Kunstleve, Paul D Adams, Martyn D Winn, Laurent C Storoni, and Randy J Read. Phaser crystallographic software. Journal of applied crystallography, 40(4):658–674, 2007.

[54] Dorothee Liebschner, Pavel V Afonine, Matthew L Baker, Gábor Bunkóczi, Vincent B Chen, Tristan I Croll, Bradley Hintze, L-W Hung, Swati Jain, Airlie J McCoy, et al. Macromolecular structure determination using x-rays, neutrons and electrons: recent developments in phenix. Acta Crystallographica Section D: Structural Biology, 75(10):861–877, 2019.

[55] Pavel V Afonine, Ralf W Grosse-Kunstleve, Nathaniel Echols, Jeffrey J Headd, Nigel W Moriarty, Marat Mustyakimov, Thomas C Terwilliger, Alexandre Urzhumtsev, Peter H Zwart, and Paul D Adams. Towards automated crystallographic structure refinement with phenix. refine. Acta Crystallographica Section D: Biological Crystallography, 68(4):352–367, 2012.

[56] Paul Emsley and Kevin Cowtan. Coot: model-building tools for molecular graphics. Acta crystallographica section D: biological crystallography, 60(12):2126–2132, 2004.

[57] Warren L DeLano. The pymol molecular graphics system. http://www.pymol. org/, 2002.

[58] Sunhwan Jo, Taehoon Kim, Vidyashankara G Iyer, and Wonpil Im. Charmm-gui: a web-based graphical user interface for charmm. Journal of computational chemistry, 29(11):1859–1865, 2008.

[59] Jing Huang, Sarah Rauscher, Grzegorz Nawrocki, Ting Ran, Michael Feig, Bert L De Groot, Helmut Grubmüller, and Alexander D MacKerell Jr. Charmm36m: an improved force field for folded and intrinsically disordered proteins. Nature methods, 14(1):71–73, 2017.

[60] David Van Der Spoel, Erik Lindahl, Berk Hess, Gerrit Groenhof, Alan E Mark, and Herman JC Berendsen. Gromacs: fast, flexible, and free. Journal of computational chemistry, 26(16):1701–1718, 2005.

[61] Shuichi Miyamoto and Peter A Kollman. Settle: An analytical version of the shake and rattle algorithm for rigid water models. Journal of computational chemistry, 13(8):952–962, 1992.

[62] Kenno Vanommeslaeghe and Alexander D MacKerell Jr. Automation of the charmm general force field (cgenff) i: bond perception and atom typing. Journal of chemical information and modeling, 52(12):3144–3154, 2012.

[63] Robert T McGibbon, Kyle A Beauchamp, Matthew P Harrigan, Christoph Klein, Jason M Swails, Carlos X Hernández, Christian R Schwantes, Lee-Ping Wang, Thomas J Lane, and Vijay S Pande. Mdtraj: a modern open library for the analysis of molecular dynamics trajectories. Biophysical journal, 109(8):1528–1532, 2015.

[64] Sophie S Spurr, Elliott D Bayle, Wenyu Yu, Fengling Li, Wolfram Tempel, Masoud Vedadi, Matthieu Schapira, and Paul V Fish. New small molecule inhibitors of histone methyl transferase dot1l with a nitrile as a nontraditional replacement for heavy halogen atoms. Bioorganic & Medicinal Chemistry Letters, 26(18):4518–4522, 2016.

